# Dual states of Bmi1-expressing intestinal stem cells drive epithelial development and tissue regeneration

**DOI:** 10.1101/2022.09.30.509798

**Authors:** Nicholas R. Smith, Sidharth K. Sengupta, Patrick Conley, Nicole R. Giske, Christopher Klocke, Brett Walker, Noelle McPhail, John R. Swain, Yeon Jung Yoo, Ashley Anderson, Paige S. Davies, Nasim Sanati, Theresa N. Nguyen, Kristof Torkenczy, Andrew C. Adey, Jared M. Fischer, Guanming Wu, Melissa H. Wong

## Abstract

Intestinal development, response to injury and disease states rely upon balanced stem cell proliferation. Historically, two subtypes of intestinal epithelial stem cells (ISCs)—slow-cycling/label-retaining, and actively-cycling/canonical Wnt-dependent—coordinate to drive proliferation and regulate epithelial renewal during adult tissue homeostasis and injury response. Recent studies focused on Bmi1-expressing cells revealed that differentiated Bmi1^+^ enteroendocrine cells could dedifferentiate towards a canonical Wnt-dependent stem cell state, calling into question the dogma that a dual stem cell axis regulates epithelial proliferation. Herein, we identify stem cell function in a Bmi1^+^ cell population in early murine intestinal development prior to the establishment of canonical Wnt-dependent, Lgr5-expressing ISCs. In-depth analyses of developmental Bmi1^+^ ISCs using lineage-tracing and single cell RNA-sequencing reveal their distinct identity and capacity to differentiate into Lgr5^+^ ISCs and other differentiated lineages. Further, during *in utero* development, the Bmi1^+^ ISCs initially exists in a highly proliferative state then transitions to a slow-cycling state, with the emergence of actively-cycling Lgr5^+^ ISCs. In adult tissue, Bmi1^+^ ISCs are a distinct population that re-express developmental gene and protein profiles, and a non-canonical Wnt signaling signature in response to injury and in human colorectal tumors. Further, developmental Bmi1^+^ ISCs are distinct from Lgr5^+^ ISCs and the previously identified differentiated Bmi1^+^ progenitor cells. Re-evaluation of an under-appreciated Bmi1^+^ ISC population with fundamental importance in intestinal development re-establishes the importance of the dynamic interplay between discrete ISC populations that are regulated by opposing Wnt signaling pathways. This finding opens opportunities and targetable pathways to augment regeneration or inhibit tumorigenesis.

## Introduction

Coordinated regulation of intestinal stem cell (ISC) proliferation drives continual epithelial renewal underlying development, homeostasis, tissue regeneration, and disease. In the mature intestine, active-proliferating ISCs expressing the canonical Wnt target gene, Lgr5 (<u>L</u>eucine-rich repeat-containing <u>G</u>-protein coupled <u>r</u>eceptor-5)^1,2^ is responsible for the rapid renewal of the epithelial cell layer. A second stem cell population expressing the chromatin regulator, Bmi1 (B lymphoma Mo-MLV insertion)^3^ represents a slow-cycling ISC population within the intestinal stem cell niche^4,5^. Data supports the dogma that together these ISC populations drive the proliferation-to-differentiation gradient to support epithelial lineage differentiation (absorptive and secretory cells^1,6–8^), and protect against accumulated injury^9,10^.

Bmi1^+^ intestinal epithelial cells are heterogeneous, comprised mainly of terminally differentiated enteroendocrine (EE) cells (~90%)^11–13^, and a small subpopulation of non-EE cells localized to the crypt-base^12^. Historically, Bmi1^+^ ISCs were considered a “reserve” ISC due to their +4 position in the crypt, label retention^3,14^, capacity for lineage tracing^12,15,16^, and stem cell identity in other organs^5,17–19^. Despite these robust data, whether or not Bmi1^+^ cells harbor ISC capacity remains controversial.

Evidence for cellular plasticity in differentiated cells^11,13,20,21^ to give rise to stem populations in response to injury presents alternative mechanisms for tissue regeneration. Bulk RNA-sequencing data support the reversion of differentiated Bmi1^+^EE cells into secretory or epithelial progenitor cells in response to radiation injury^11,12^. The designated progenitor cells gain characteristics and function of Lgr5^+^ ISCs through the canonical Wnt target gene, Achaete-scute family bHLH transcription factor 2, *Ascl2*^13^. However, these datasets^11,13^ lack adequate single cell resolution of the heterogeneous Bmi1^+^ cell population, and thus precludes definitive evidence to exclude the possibility that a subpopulation of Bmi1^+^ cells function as ISC in the intestine.

Whether Bmi1^+^ cells independently harbor meaningful stem cell capacity and biologically contribute to intestinal biology continues to be downplayed^22^. Herein, we demonstrate that Bmi1^+^ cells represent the predominant epithelial ISC in early intestinal development and precede existence of canonical Wnt-dependent Lgr5^+^ ISCs. We reveal that early in development, Bmi1^+^ ISCs are highly proliferative, then transition to a slow-cycling state with the emergence of Lgr5^+^ ISCs. Combining single cell RNA-sequencing (scRNA-seq) with bulk population ATAC-seq (Assay for Transposase Accessible Chromatin sequencing) analyses, we identify that a non-canonical Wnt signaling pathway correlates with the temporal presence of the highly proliferative developmental Bmi1^+^ ISC population. scRNA-seq analyses reveal a discrete subpopulation among adult homeostatic Bmi1^+^ cells that harbors stem cell gene signatures, as well as a proliferative, development-like Bmi1^+^ ISC subpopulation that emerges in response to injury and in tumorigenesis. Our findings identify different intestinal stem cell states in Bmi1^+^ epithelial cells in the context of development, adult homeostasis, regeneration and tumorigenesis, highlighting the diverse contribution of Bmi1^+^ ISC to intestinal epithelia maintenance.

## Results

### Adult homeostatic Bmi1-GFP^+^ cells revert to a proliferative developmental-like state in culture

Recent studies downplay a stem cell role for subsets of Bmi1^+^ epithelial cells^11–13,22^, yet isolated Bmi1^+^ cells grow undifferentiated spheroids in culture^23^. To investigate the Bmi1^+^ subpopulation with progenitor or stem cell function, we FACS-isolated Bmi1-GFP^+^ epithelium from adult murine intestines (Fig S1A) and subjected cells to standard growth conditions^24^. Single Bmi1-GFP^+^ cells grew into highly proliferative, multi-cellular spheroid structures distinct from Lgr5^+^-derived, budded organoids^23,24^ (Fig 1A, top panel), which we previously demonstrated grow in a R-spo1-independent fashion^23^. Unlike structures grown from single Lgr5^+^ ISCs, Bmi1-derived spheroids were comprised almost entirely of Bmi1-GFP^+^ cells and were highly proliferative, differing from their *in vivo* counterparts localized to the base of intestinal crypts, and seldomly dividing (Figure 1A, top panel). This spheroid growth phenotype is identical to reported findings from developing intestinal epithelia^25,26^, and from Apc^Min^ adenomas^27^. Given these similarities, we hypothesized a developmental role for Bmi1^+^ cells. In E14.5 intestines from Bmi1-reporter mice^7^, Bmi1-GFP is expressed throughout the intestinal epithelium, stroma, and muscle layer (Fig 1A). We FACS-isolated EpCAM^+^/GFP^+^ cells from E14.5 Bmi1-GFP intestines (Fig S1B) and cultured them under standard conditions. Similar to the adult population, E14.5 intestinal epithelial cells produced large spheroids that ubiquitously expressed Bmi1-GFP and could be serial passaged (i.e., self-renew, Fig 1A). This indicated that adult Bmi1-GFP cells contain a subpopulation with self-renewal capacity when stimulated *ex vivo*.

**Figure 1.**
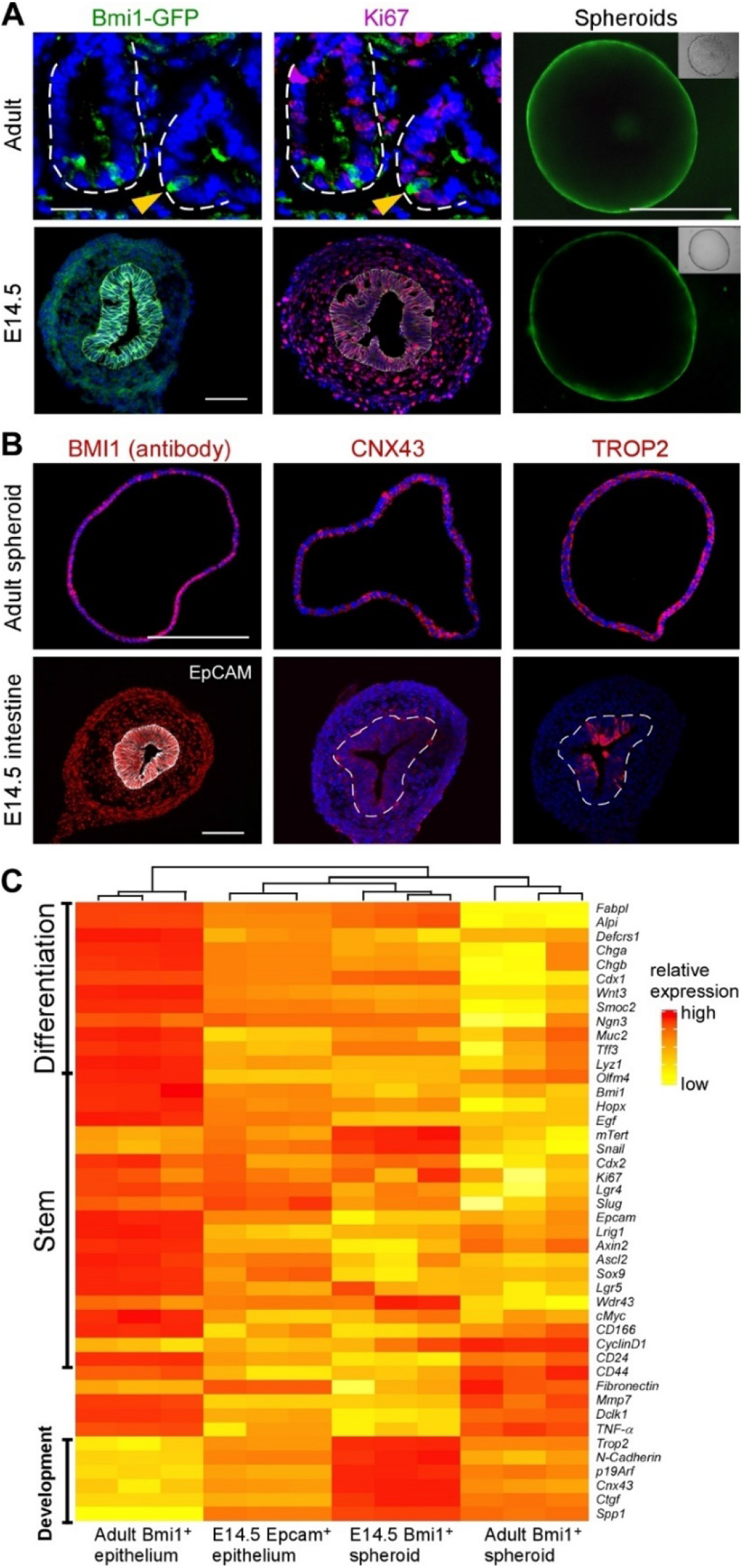
*Ex vivo* cultured adult murine Bmi1-GFP^+^ cells revert to a developmental-like state. (A) Sections of adult small intestine (top panel) and embryonic (E)14.5 developing intestine (bottom panel) from Bmi1-GFP reporter mice stained with antibodies against GFP (left, green) and Ki67 (middle, red). (Right) FACS-isolated Bmi1-GFP^+^ cells from adult (top) and E14.5 developing intestines (bottom) generate large Bmi1-GFP-expressing spheroid structures *ex vivo*. Inset: phase contrast image. (B) Sections of *ex vivo* adult Bmi1-GFP spheroids (top panel) and E14.5 intestine (bottom panel) stained with antibodies against Bmi1 (left, red), Cnx43 (middle, red) and Trop2 (right, red). Scale bar = 50 μm. (C) Heatmap of qRT-PCR gene expression from isolated Adult Bmi1-GFP^+^ intestine, E14.5 EpCAM^+^ intestine, E14.5 Bmi1^+^ spheroids and Adult Bmi1-GFP^+^ spheroids. Data are presented as ΔΔCT relative to Gapdh and normalized to E17.5 Epcam^+^ cells. Data are representative of N=3 independent experiments from at least N=6 total mice.

To identify similarities, we stained adult Bmi1^+^ spheroids, and E14.5 intestine with antibodies against Bmi1. Nuclear Bmi1 staining was consistent with Bmi1-reporter expression, observed in spheroids and developing EpCAM^+^ cells (Fig 1B). To establish that adult spheroids exist in a developmental state, we stained spheroid sections for developmentally expressed proteins, and identified connexin 43 (CNX43) and tumor-associated calcium signaling transducer 2 (TROP2)^26^ in adult Bmi1-spheroids and the developing intestine (Fig 1B), but not in homeostatic adult epithelia (see Fig 6C). We next FACS-isolated Bmi1^+^ epithelia from adult intestine, adult Bmi1^+^ spheroids, E14.5 intestine, and E14.5 Bmi1^+^ spheroids, then compared their gene expression profiles. Adult Bmi1-GFP^+^ cells expressed high levels of differentiated genes, including EE genes *ChgA* and *ChgB*^28,29^, whereas adult spheroids had lower expression of these genes as well as other differentiation-associated genes (e.g. *Alpi, Muc2, Lyz1*) and upregulated expression of developmental genes (e.g. *Trop2, Cnx43*), displaying similar expression to developmental tissue (Fig 1C). These data indicate that *ex vivo* conditions drive adult Bmi1-GFP^+^ cells toward a developmental state. Moreover, adult Bmi1-GFP^+^cells generate spheroid structures rather than phenotypic Lgr5-budded organoid structures, providing evidence that a subpopulation of adult Bmi1-GFP^+^ cells are progenitors or ISCs that are discrete from Lgr5 ISCs.

### Bmi1-GFP^+^ cells are the major epithelial population in the developing intestine

Clear proliferative differences in adult and developmental Bmi1^+^ intestinal epithelium (i.e. Ki67, Figure 1A) implied that developmental Bmi1^+^ cells may harbor stem or progenitor attributes. Therefore, we investigated Bmi1^+^ cells in the developing intestine from E12.5 to E17.5 and compared them to Lgr5^+^ ISCs during the same timeframe. This period of development spans changes in epithelial proliferation and morphology from pseudo-stratified to a single columnar layer that precedes crypt-villus formation, and the residence of Lgr5^+^ ISC within the crypt region. We compared Bmi1^+^ and Lgr5^+^populations using GFP reporter mice to examine the developmental relationship between these discrete cellular pools. The E12.5 EpCAM^+^ intestinal epithelia predominantly expressed Bmi1-GFP identified by immunofluorescence, relative to non-GFP, control mice (C57Bl/6, Figure 2A) and by FACS analyses of dissociated cells (Figure 2B and S2). In contrast, at this timepoint, Lgr5-GFP was expressed by only a minor subset of intestinal epithelia (Figure 2B, C, S2, 3.8%) consistent with published data^30–32^. FACS analyses of additional developmental time points (Figure S2) revealed that the number of Bmi1-GFP^+^ cells decreased as Lgr5-GFP^+^ increased (Figure 2D), with the highest percentage of Lgr5-GFP^+^ ISC relative to total epithelia at E15.5 (Figure 2E). The decrease in Bmi1-GFP^+^ cells and increase in Lgr5^+^ ISCs corresponded with reported initiation of a canonical Wnt signaling environment^31^. These data identify an inverse relationship between Bmi1^+^ and Lgr5^+^ populations across intestinal developmental, and that the canonical Wnt signaling pathway may be involved in shifting the ratio between these populations.

**Figure 2.**
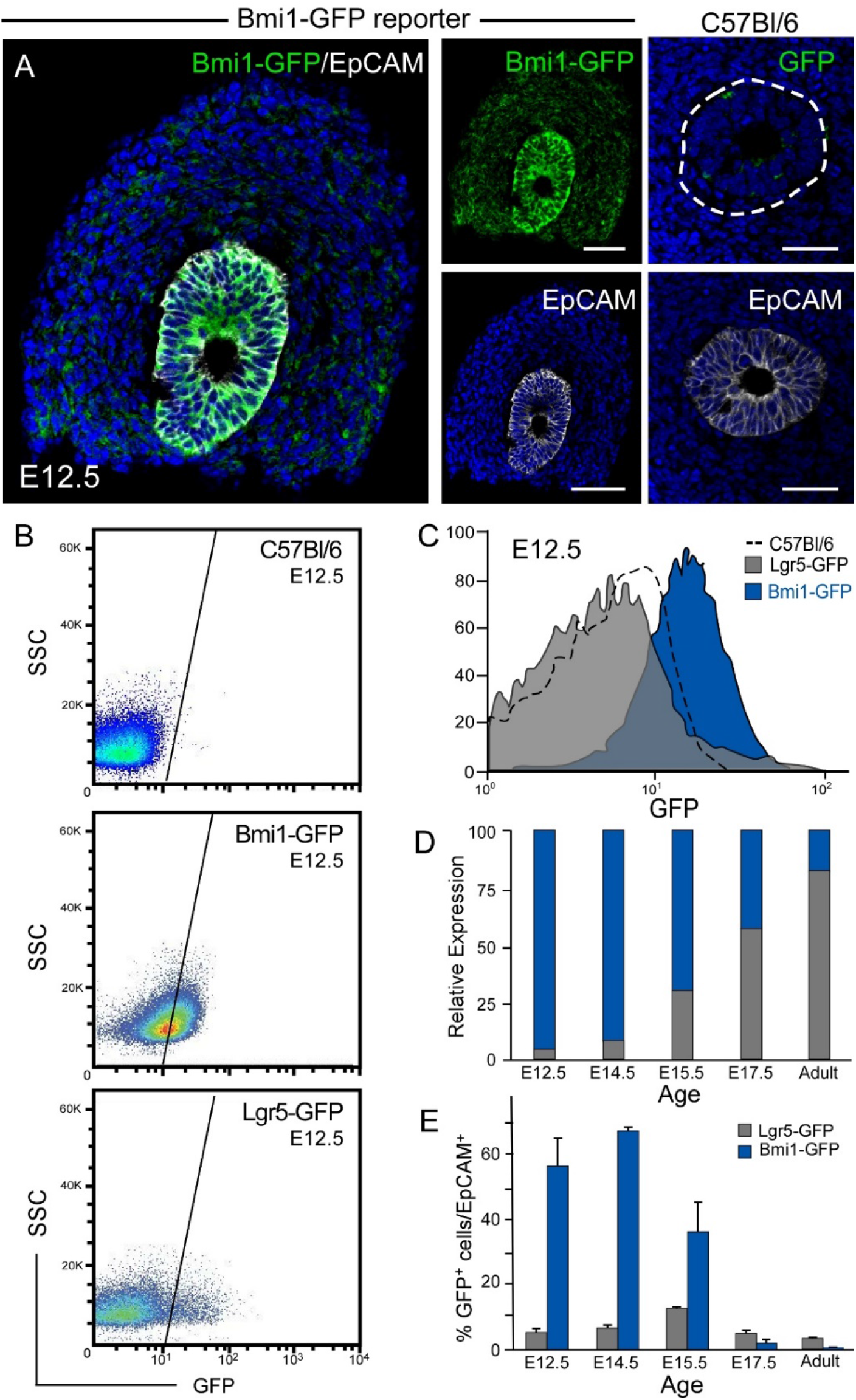
Bmi1-GFP is highly expressed during early murine intestinal development. (A) Tissue sections from embryonic day (E)12.5 Bmi1-GFP^+^ or C57Bl/6 WT control mice stained with antibodies against GFP (green) and EpCAM (white). Dashed line denotes epithelial-stromal boundary. Scale bar = 25 μm. (B) FACS plots of epithelial (EpCAM^+^) cells from E12.5 developing intestines from C57Bl/6 (top), Bmi1-GFP^+^ (middle) and Lgr5-GFP^+^ (bottom) intestines. (C) Histogram of GFP signal from E12.5 FACS plots comparing C57Bl/6 (dashed line), Bmi1-GFP (blue) and Lgr5-GFP (gray). (D) Bar graph depicting the relative expression of Bmi1-GFP^+^ and Lgr5-GFP^+^ epithelial cells in intestines across developmental time and in adult intestine, analyzed by flow cytometry (i.e. ratio of Bmi1^+^:Lgr5^+^cells). (E) Bar graph depicting the percent of Bmi1-GFP^+^ and Lgr5-GFP^+^ per total EpCAM^+^ cells across developmental time and in adult tissues, analyzed by flow cytometry (i.e. Bmi1^+^ or Lgr5^+^ cells relative to total epithelial cells). Data are representative of at least N=3 independent experiments from N=6 mice per genotype.

### Developmental Bmi1^+^ epithelial cells give rise Lgr5-GFP^+^ ISCs and differentiated lineages

Temporal evaluation of Bmi1^+^ and Lgr5^+^ populations indicate that Bmi1^+^ cells predominate within the epithelium during early development (Figure 2) indicating that Bmi1^+^ cells precede, and therefore may give rise to the Lgr5^+^ ISC population. To test this hypothesis, we first explored the lineage tracing capacity of Bmi1 and Lgr5 cells at early developmental time points. We induced lineage tracing from Bmi1-Cre^ERT^ and Lgr5-GFP-IRES-Cre (Lgr5Cre^ERT^) mice crossed onto a TdTomato (TdT) reporter as early as E8.5, at a timepoint previously shown to have minimal Lgr5 expression within the intestine^32^. We detected robust lineage tracing 9 days post-induction (E17.5) within the intestinal epithelium of Bmi1-Cre^ERT^/TdT mice, whereas very few TdT lineage marked cells were detected in Lgr5-Cre^ERT^/TdT embryos (Figure 3A). Notably, some Bmi1-derived epithelial lineages were localized within the intervillus region, the location of proliferative progenitors and Lgr5-GFP^+^ cells at E17.5 (Figure 3A, white arrowheads)^33,34^. Importantly, lineage tracing induced at E8.5, E9.5 (not shown) or E10.5 (Figure S3B) produced identical results from the Bmi1 promoter, indicating that lineage tracing from E8.5 was not a result of endodermal progenitor cell labeling.

**Figure 3.**
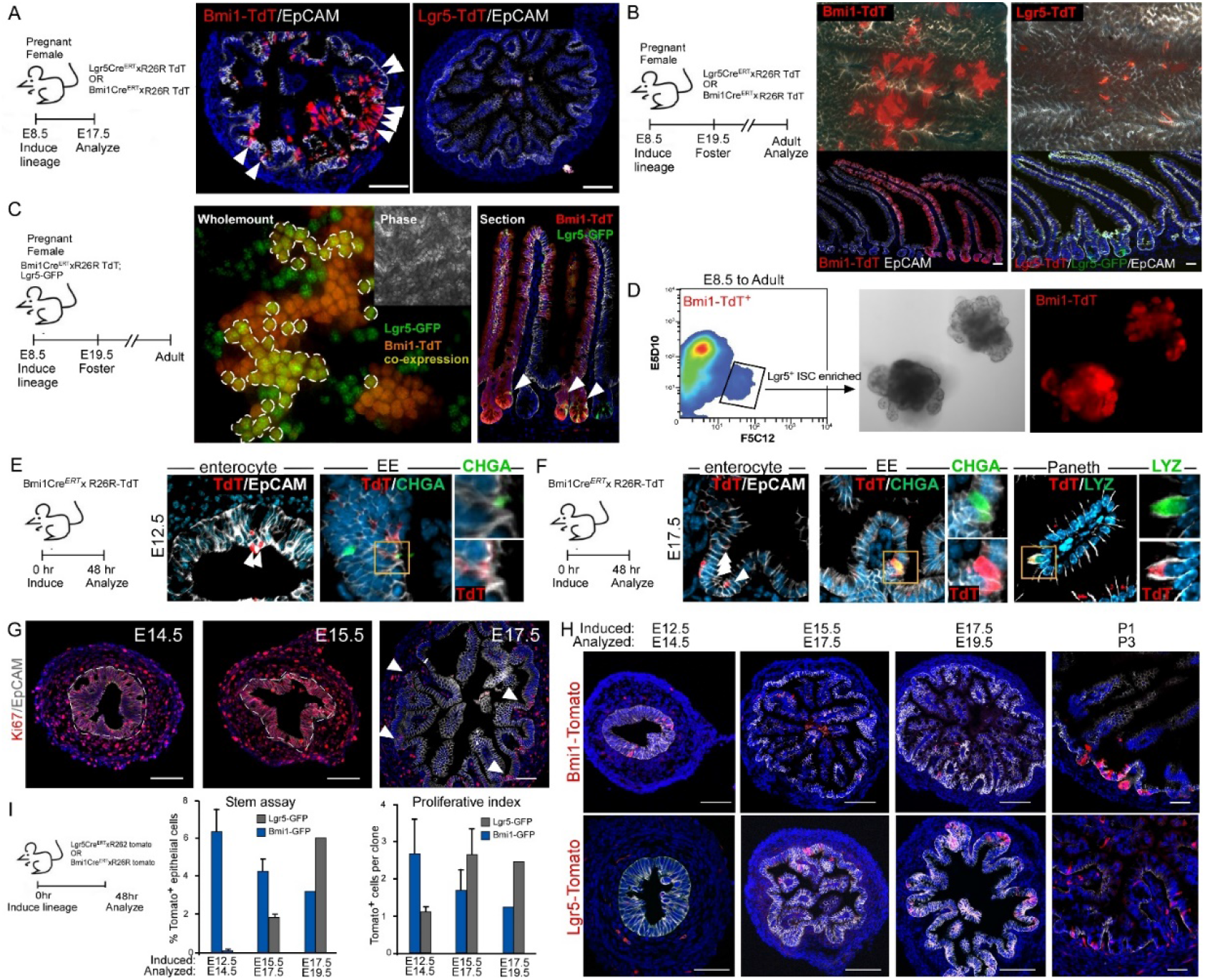
Bmi1-expressing intestinal epithelial cells in early development give rise to adult Lgr5-GFP^+^ ISCs. (A) Left: Experimental paradigm for lineage tracing from Bmi1- or Lgr5-expressing cell populations at E8.5 with analyses at E17.5. Right: Intestinal tissues from Bmi1 or Lgr5 traced tissues with lineage-derived cells marked by TdTomato (TdT) expression (red) and co-stained for EpCAM (white), scale bar = 25 μm. (B) Left: Experimental paradigm for lineage tracing from Bmi1- or Lgr5-expressing cells at E8.5 and fostering animals for adult tissue analysis. Top Right: Adult wholemount intestine images, and bottom right: adult tissue sections from Bmi1- or Lgr5-traced tissues showing lineage^+^ cells marked by TdT (red), scale bar = 25 μm. Tissue sections: TdT (red), EpCAM (white) and Lgr5-GFP (green), scale bar = 25 μm. (C) Left: Experimental paradigm for E8.5 lineage tracing from Bmi1^+^ cells in an Lgr5-GFP reporter background, followed by analyses in adult animals. Right: Wholemount intestine image of crypt bottoms showing Bmi1-derived cell lineage marked by TdT (red) and Lgr5-GFP^+^ ISCs (green). Co-expressed crypts are marked by white dashed outlines. Phase contrast image inset, scale bar = 25 μm. Tissue sections showing Bmi1-derived cell lineage with TdT (red), stained for EpCAM (white) and Lgr5-GFP (green). Arrowheads denote TdT^+^ crypts with Lgr5-GFP^+^ ISCs, scale bar = 25 μm. (D) Left: FACS plot of adult Bmi1-lineage traced cells stained with antibodies E5D10 and F5C12 to isolate the Lgr5-enriched E5D10^lo^ F5C12^hi^ population^23^. This cell population gives rise to TdT-marked intestinal enteroids in *ex vivo* cultures, phenotypically identical to Lgr5-derived enteroids. Data are representative of N=7 mice from at least 3 independent experiments. (E) Short term, 48 hr lineage tracing from the *Bmi1* promoter from E12.5 give rise to enterocytes (expressing EpCAM), and enteroendocrine (EE) cells (expressing Chromagranin A, CHGA). (F) Short term, 48 hr lineage tracing from the *Bmi1* promoter from E17.5 give rise to enterocytes (expressing EpCAM), EE cells (expressing CHGA), Paneth precursors (expressing lysozyme, LYZ) and tuft cells (expressing doublecortin like kinase 1, DCLK1). (G) Intestinal tissue sections from E14.5, E15.5 and E17.5 intestines stained with antibodies against Ki67 (red) and EpCAM (white). White arrows denote regions of proliferating cells localized to intervillus zone, scale bar = 25 μm. (H) Left: Experimental paradigm for 48 hr lineage trace from Bmi1- or Lgr5-expressing cells at E12.5, E15.5 or E17.5 and analyzed 2 days later. Right: Bar graph quantification of percent EpCAM^+^ cells lineage marked by TdT and size of TdT^+^ clones for Bmi1 (blue bars) and Lgr5 (gray bars) across developmental time. Data are presented as mean +/− SEM. (I) Intestinal tissue sections from Bmi1- or Lgr5-lineage tracing with lineage-derived cells marked by TdT (red) and stained for EpCAM (white). Scale bar = 25 μm. Data are representative of N=3-30 mice from at least 2-6 independent experiments.

Lgr5^+^ ISCs localize to the inter-villus region during development^34^ where Bmi1-TdT^+^ cells resided, and thus we explored the possibility that developmental Bmi1^+^ cells could give rise to Lgr5^+^ ISCs. We repeated lineage tracing in Bmi1-Cre^ERT^/TdT mice induced at E8.5 but analyzed the intestinal epithelium for TdT^+^ clones in adult animals aged to ~8 weeks. Consistent with the E17.5 analyses, large TdT^+^ clones were detected along the length of the small intestine and within the colon (Figure 3B, Figure S3A) from Bmi1-Cre^ERT^ animals by wholemount and tissue section analyses but were nearly absent from Lgr5-Cre^ERT^/TdT intestines. Bmi1-TdT^+^ clones appeared to encompass clusters containing multiple labeled adjacent crypts, as expected if tracing was from an early developmental progenitor or stem cell population.

To further investigate this, we evaluated E8.5 induced Bmi1-lineage traced cells in E17.5 (Figure S3C) and adult animals (Figure 3C) that had both Bmi1-Cre^ERT^ TdT and the Lgr5 reporter expression (i.e. Lgr5-GFP). In this paradigm, Bmi1 progeny marked by TdT that gave rise to Lgr5^+^ cells would co-express TdT and GFP. As demonstrated in Figure 3A-B and Figure S3C, induction of Cre expression at E8.5 drives lineage tracing from the Bmi1 and not the Lgr5 promoter, and thus the majority of TdT^+^ cells were Bmi1-derived. In adult tissue, entire crypt and adjacent villus TdT^+^epithelia overlap with Lgr5-GFP-expression within crypts (Figure 3C), whereas in E17.5 co-expression of TdT (Bmi1-derived lineage) and GFP (Lgr5 reporter) is appreciable (Figure S3C). Additionally, we evaluated Bmi1-lineage for functional Lgr5 stem cell capacity, using approaches that do not rely upon the Lgr5-reporter mouse. To do this, we dissociated epithelium from lineage traced Bmi1-Cre^ERT^/TdT adult intestines then stained the cells with the monoclonal antibody F5C12 that enriches for the Lgr5-ISC population^23^. Bmi1-TdT^+^ epithelial cells with co-expression of F5C12^+^ were isolated by FACS and displayed robust growth under *ex vivo* enteroid culture conditions^24^ that were phenotypically identical to Lgr5-derived organoids with budded crypt-like structures (Figure 3D). Resultant enteroids were Bmi1-TdT^+^ and expanded through multiple passages, indicating that developmental Bmi1^+^ ISCs have the capacity to generate adult Lgr5^+^ ISCs.

To determine if developmental Bmi1^+^ cells can give rise to downstream differentiated lineages, we analyzed the protein expression in Bmi1-TdT^+^ cells after a 48 hr trace initiated at E12.5 or E17.5 (Figure 3E, F). In both time points, lineage^+^ cells were EpCAM^+^, indicating that Bmi1^+^ cells divide and generate differentiated enterocytes. In E12.5-to-E14.5 lineage analyses CHGA^+^ EE were extremely rare, but we observed lineage tracing to EE lineages. Further, while Paneth cell progenitors are detected in the intervillus region, fully differentiated Paneth cells that are LYZ^+^ are not yet readily detected in E14.5 intestines^35^, and thus we were unable to detect an adequate number of LYZ^+^ cells to definitively assess Bmi1^+^ ISC lineage relationships. Similarly, in E17.5-to-E19.5 intestines (lineage traced from E17.5), rare EE (CHGA^+^), and Paneth (LYZ^+^) cells co-expressed TdT (Figure 3F), while tuft cell detection^36^ fell below the level for analyses of lineage. These data indicate that Bmi1^+^ cells can give rise to differentiated epithelial lineages (i.e. enterocytes, enteroendocrine and Paneth cells), thus supporting that early developmental Bmi1^+^ cells harbor hallmark stem cell characteristics, self-renewal capacity (Figure 1A) and the ability to give rise to downstream lineages (Figure 3E,F). Of impactful significance, we also demonstrated that the developmental Bmi1^+^ ISC population reside upstream of the Lgr5^+^ ISC population.

### Highly proliferative developmental Bmi1^+^ ISCs transition to a slow-cycling state coincident with the emergence of Lgr5^+^ ISCs

Given that developmental Bmi1^+^ ISCs are highly proliferative but adult intestinal homeostatic Bmi1-expressing cells are slow-cycling, we explored the timing and of this proliferative state transition by evaluating the proliferative status of Bmi1^+^ ISC across developmental timepoints. Consistent with previous reports, Ki67^+^ proliferative cells represent the majority of the intestinal epithelium in early development (E14.5), then become restricted to the intervillus zone in late development (E17.5; Figure 3G, white arrowheads). To determine if reduced proliferation was due to a decrease in ISC numbers or rate of proliferation, we set out to evaluate proliferative capacity and clonal expansion of Bmi1 and Lgr5 cells by utilizing short-term (48 hr) lineage tracing in Bmi1-Cre^ERT^/TdT and Lgr5-Cre^ERT^/TdT mice at three developmental timepoints, E12.5, 15.5 and 17.5. Consistent with GFP reporter expression (Figure 2E), at E12.5, Bmi1^+^ ISC numbers were highest with ~6% of EpCAM^+^/Bmi1^+^ cells giving rise to downstream lineages by E14.5, while <0.1% of EpCAM^+^/Lgr5^+^ ISCs generated labeled progeny (Figure 3H, I). Determining number of cells within a TdT^+^ clone provided a proliferative index of discrete ISC populations. We found that Bmi1-lineage traced clones averaged 2-3 cells in size, indicating at least one cell division during the 48 hr period, while rare Lgr5-TdT^+^ clones were consistently only 1 cell in size—thus not dividing during this analytic period (Figure 3H, Figure S4).

We found that Bmi1-initiated lineage tracing was reduced to ~4 and 3% of EpCAM^+^ cells from E15.5 and E17.5 induction, while Lgr5-derived tracing was significantly increased relative to E12.5, with ~2 and 6% EpCAM^+^ cells (Figure 3H and Figure S4). Further, analysis of TdT^+^ clone size indicated that Bmi1^+^ ISCs had diminished proliferative capacity (1 cell per clone at E17.5), while the Lgr5-TdT clones were larger (3 cells per clone), indicating significant proliferation of the Lgr5^+^ population in late development (Figure 3H and Figure S4). These data indicate the reduction in Bmi1^+^ ISC proliferation occurs during mid-development and coincides with the emergence of the canonical Wnt-responsive and proliferative Lgr5^+^ ISC population.

### Developmental Bmi1^+^ ISCs from E12.5 and E17.5 intestines are distinct entities

Given the increasing level of cellular heterogeneity that occurs as the intestine develops, analyses of bulk cellular populations lack adequate resolution to appreciate rare stem populations and their downstream progeny. Therefore, to compare defining features of Bmi1^+^ ISCs and their downstream Bmi1-expressing progeny in E12.5 and E17.5 intestines, we surveyed >16,000 murine Bmi1-GFP^+^ epithelial cells through droplet-based 3’ scRNA-seq. After filtering out low-quality cells, normalization, dimensionality reduction, and clustering analyses, cells were merged together after batch correction using the Harmony approach^37^ to facilitate direct comparison of E12.5 and E17.5 populations, and then visualized after projected by Uniform Manifold Approximation and Projection (UMAP)^38^ (Figure 4A). There was 10% overlap between E12.5 and E17.5 Bmi1^+^ cells indicating that these populations are fundamentally distinct. Thus, we analyzed datasets independently at each time point to assess stem and differentiating populations. For the E12.5 Bmi1^+^ population, although the cells were predominantly homogeneous, we identified five distinct subpopulations (Figure 4B) with unique characteristics, four with characteristic undifferentiated cells and a fifth with EE gene expression (see Table S1). Marker analyses revealed that all four undifferentiated clusters differentially expressed proliferative genes (e.g. *Cdk1, Ccnd1, Bub3, Plk1*) and factors associated with stem or progenitor states, including the pluripotency factor *Phf5a* in cluster 0, numerous Hox genes and other homeobox genes—pioneering transcription factors that are important for chromatin-mediated gene expression and maintenance of stem cell state^39,40^ in cluster 1 and 2, genes that inhibit lineage differentiation (e.g. *Dlk1, Notch 1, Id3^41,42^*) in cluster 2, and key developmental gene that regulate EMT or cell fate decisions (e.g. *Tead2, Sox4*) in clusters 2 and 3. We used Cellular Trajectory Reconstruction Analysis using gene Counts and Expression (CytoTRACE)^43^ analyses, a computational approach to infer the differential states of individual cells based on the significantly strong correlation between the number of expressed genes and differential states, and pseudotime analyses and identified cluster 0 as the most undifferentiated population of E12.5 Bmi1^+^ cells (Figure 4C, Figure S5B,C). The small cluster 4 representing 11 cells harbored DEGs involved in EE lineage specification (e.g. *Chga, Chgb, Arx, Neurod1, Isl1, Chga* ^28,44–47^), despite only very rare E12.5 Bmi1^+^ cells expressing EE associated proteins (Figure 4B).

**Figure 4.**
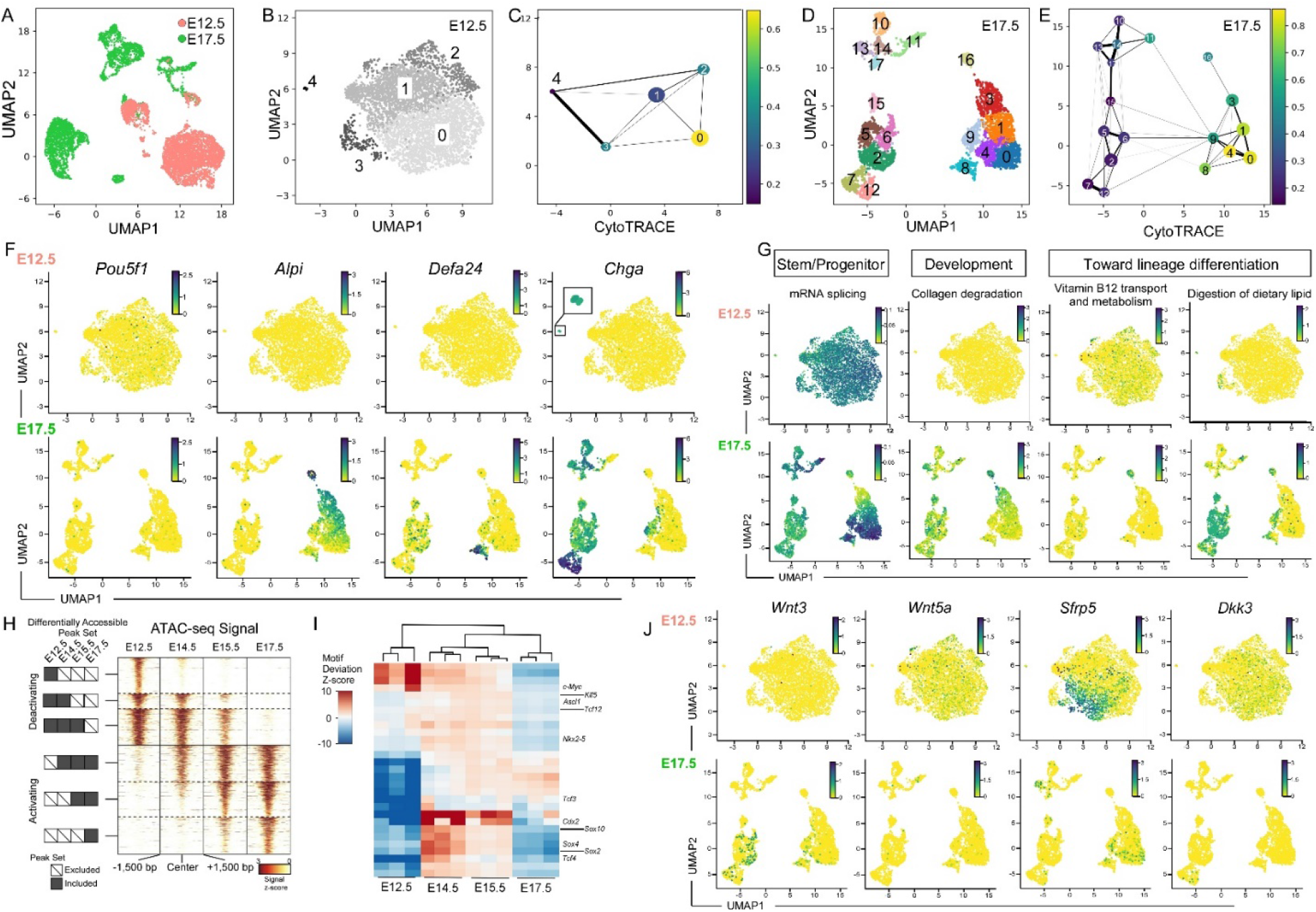
Bmi1-expressing ISCs represent discrete entities. (A) UMAP plot of E12.5 and 17.5 Bmi1^+^ scRNA-seq datasets integrated using the Harmony approach. (B-E) UMAP plots and cytoTRACE analyses of E12.5 dataset (B,C) and E17.5 (D,E). (F) UMAP plots of Bmi1-GFP^+^ cells from E12.5 (top) and E17.5 (bottom) displaying lineage-specific expression for pluripotency transcription factor, *Pou5f1* (Oct4), and genes expressed in differentiated enterocytes: alkaline phosphatase (*Alpi);* Paneth cells: α-defensin 24 precursor (*Defa24);* and enteroendocrine cells: Chromogranin A (*Chga*). A small 11 cell cluster of cells in the E12.5 *Chga* UMAP is boxed and enlarged as an inset. (G) UMAP plots of Bmi1-GFP^+^ cells from E12.5 (top) and E17.5 (bottom) depicting activities of pathways in Stem/Progenitor: mRNA splicing; Developmental: collagen degradation; and Differentiation: vitamin B12 transport and metabolism, and digestion of dietary lipid. (H) Heatmap of activating and deactivating ATAC-seq signal (±1.5kb window from peak center) across different developmental timepoints of Epcam^+^ cells. (I) Hierarchical clustering of known motif HOMER enrichment analysis from ATAC-seq data of developmental EpCAM^+^ cells. (J) Bmi1-GFP^+^ cells from E12.5 (top) and E17.5 (bottom) validating ATAC-seq Wnt signaling pathway results. Canonical Wnt ligand (*Wnt3*) and non-canonical ligand (*Wnt5a*) gene expression and inhibitors of the canonical Wnt pathway, secreted frizzle related protein 5 (*Sfrp5*) and Dickkoft WNT signaling pathway inhibitor 3 (*Dkk3*) are shown.

In contrast to the predominantly homogeneous cell populations at E12.5, E17.5 Bmi1-expressing cells clustered into 17 discrete subpopulations representing stem/progenitor features, enterocyte, Paneth, and EE cell identity (Figure 4D, Figure S5E, Table S1). Clusters 0 and 4 were the most undifferentiated populations with multiple stem cell-associated DEGs, including *Tead2, Shh*, and *Cldn6*. CytoTRACE (Figure 4E) and pseudotime inference analyses (Figure S5D, E), along with DEGs of clusters, revealed lineage differentiation down three pathways: enterocytes, Paneth and EE cells (Figure S5E, Table S1). Specifically, DEGs in clusters 0-4-1-3-16 reflected increasing differentiation status along the enterocytic lineage^45^, with similar trends for the Paneth cell lineage, and the EE lineage^44^. To further explore differences across the two time points, we evaluated gene expression discrete to epithelial lineages identified in Figure 4F. As expected, *Pou5f1* (or Oct4), a pioneer transcription factor expressed in early development and important for stem cell pluripotency^48^, was highly expressed almost exclusively in E12.5 populations, whereas the alkaline phosphatase gene (*Alpi*),characteristic of mature enterocytes was not expressed in E12.5 cells, and instead was restricted to populations along the enterocyte differentiation gradient in the E17.5 population. Interestingly, a single cluster in E17.5 expressed α-Defensin (*Defa24*), a Paneth cell related gene at high levels, potentially indicating that Bmi1^+^ ISC can initiate lineage differentiate toward this important stem niche cell. These cells also expressed *Cldn7* and *Muc2*, characteristic of Paneth precursor cells reported to underlie cellular plasticity in response to tissue injury^20,21,49–51^ (Table S1). Finally, chromogranin A (*Chga*), a gene characteristic of mature EE cells^11,12,52^ was highly expressed in a number of E17.5 subpopulations, highlighting a potential role for Bmi1 in this lineage differentiation. Of note, *Chga* was almost not detected in the majority of E12.5 Bmi1^+^ cells, and only in quite high levels in cluster 4 (composed of 11 cells; Figure 4F, inset).

AUCell^53^ pathway activity analysis of the cell populations based on the Reactome pathway knowledgebase^54^ provided a global functional view of the subpopulations at each developmental time point. Pathways involved in mRNA splicing, collagen degradation, vitamin B12 transport and metabolism and digestion of dietary lipids distinguished discrete subpopulations of cells, as well as highlighted the differences between E12.5 and E17.5 populations (Figure 4G). Further, pathways for open chromatin, transcription and translation, as well as RUNX3 mediated pathways were high in the E12.5 population, while pathways involved in dietary lipid metabolism, digestion, absorption and collagen degradation were observed in non-stem cell populations at E17.5 (see also Figure S5F). Together, these data, along with lineage tracing in Figure 3E, F, indicate that the developmental Bmi1^+^ stem cell population can give rise to EE and enterocyte lineages, as well as to a subpopulation representing a Paneth cell progenitor, during early intestinal development.

To identify potential cell signaling pathways regulating the transition of Bmi1^+^ISCs from a proliferative to slow-cycling state, or involved in lineage differentiation, we performed bulk ATAC-Seq analyses of FACS-isolated EpCAM^+^ intestinal epithelial cells across developmental timepoints. Using differential chromatin accessibility analysis we identified enrichment of multiple pathways in the E12.5 population that progressively decreased at E17.5 (Figure 4H). Using identification of motif discovery within this dataset, we found enhanced utilization of transcription factors in pathways including the Hippo signaling pathway (TEAD1, TEAD2 and TEAD4)^55^, a known pathway that regulates organ development and size^56^ (Table S2). Interestingly, the Hippo pathway is a known negative regulator of the canonical Wnt signaling pathway^57^, thus its decrease in expression is consistent with the emergence of the canonical Wnt-regulated Lgr5^+^ISC during development. When we evaluated HOMER enrichment motifs for canonical Wnt-responsive loci (Figure 4H), we identified occupancy peaking at E14.5, again consistent with temporal transition from Bmi1^+^ ISCs as the primary population to the Wnt-responsive Lgr5^+^ ISC population. Global motif enrichment for Wnt-related targets was then assessed using ChromVAR^58^, which displayed a similar trend, with the canonical Wnt pathway signature peaking at E14.5 and E15.5 (e.g., *Cdx2, Tcf4* and other labeled genes; Figure 4I). Incidentally, as the analysis was conducted on all EpCAM^+^ cells, reduction in the Wnt signature at E17.5 likely reflects the decrease in Lgr5^+^ ISC numbers relative to differentiated epithelial cells, as noted in Figure 2E.

To complement the findings from ATAC-seq analyses, we examined Wnt signaling pathway effector gene expression across the scRNA-seq developmental Bmi1^+^ datasets. The canonical Wnt ligand, Wnt3 expression was localized to subpopulations of E17.5 cells that harbored *Defa24* and/or *Chga* expression, and was not expressed in E12.5 cells. Contrary to this, Wnt5a, a non-canonical Wnt ligand expression was increased in E12.5 cells, and present only in a small subset of E17.5 cells (Figure 4J). Further, canonical Wnt antagonists, *Sfrp5* and *Dkk3*, were also highly expressed in the E12.5 cells, and less so in E17.5 cells. These expression patterns indicate differential utilization of Wnt signaling pathways, where there is reliance on non-canonical signaling regulation in early development (E12.5), then a transition to dependence on the canonical Wnt microenvironment to drive (Lgr5) stem cell proliferation and maintenance in mid development (E14.5-15.5) and into adulthood. These findings indicate that the non-canonical Wnt pathway is involved in regulating early developmental Bmi1^+^ ISC proliferation.

To directly determine the differences between E12.5 and E17.5 ISC populations, we compared DEGs from cluster 0 (E12.5, light gray) and cluster 0,4 (E17.5, dark green) and identified only 23 genes shared between the two populations (Figure S5H, Table S3). The discrete nature of the two ISC populations was further visualized when they were highlighted on the UMAP comparing E12.5 and E17.5 populations (Figure S5G). Collectively, the data presented herein identifies E12.5 and E17.5 Bmi1^+^ ISC populations as distinct entities with discrete proliferative states.

### A minor subpopulation of adult Bmi1^+^ cells harbor stem cell identity

Across development we found that Bmi1^+^ ISCs transition from an active-proliferating to a slowly-cycling population, thus we asked if Bmi1^+^ cells retained stem cell capacity in the adult intestinal epithelium during homeostasis. Published data highlights Bmi1^+^ cells as a heterogeneous population in the mature intestinal epithelia with a majority of cells harboring protein and gene expression of mature EE lineages and a minor Bmi1^+^ population lacking EE expression (Figure S6A)^12^. The cluster analysis of the adult Bmi1^+^ scRNA-seq dataset identified a small stem cell population (less than 10% of the population) with upregulated proliferative and ISC gene expression (*Mki67, Cdk1, Olfm4, Ascl2;* Figure 5A, B, D), but with a minimal *Lgr5* expression (Figure 5D). To determine if adult Bmi1-ISCs shared expression with developmental ISCs, we compared DEGs from the E12.5 cluster 0 (light gray) or the E17.5 cluster 0,4 (dark green) with the adult ISC cluster (red) (Figure 5C). We found that the adult ISC population shared only 36% of the E12.5 Bmi1-ISC population gene expression, but shared 73% of the E17.5 gene expression (Figure 5C, Table S3). Alignment of cluster-based differentially upregulated genes with known cell type markers (Figure 5A, B, Table S4) identified four distinct EE cell clusters based on co-expression of secretory products, EE-EC (*Chga, Chgb, Tph1*), EE-I (*Gcg, Cck, Ghrl*), EE-D (*Ghrl, Sst, Iapp*), and EE-K (*Gip, Isl1*)^12,44,45,52^, Tuft (*Trpm5, Sox9, Dclk1*), and smaller numbers of Paneth (Pan; *Defa17, Itln1, Defa24, Lyz1, Mmp7*). Enterocyte (Ent; *Alpi, Lct, Fabp2*)^45^ subpopulations were also detected among the Bmi1-expressing cells (Figure 5A). We also identified a broad subpopulation that expressed multi-lineage genes and that did not meet the criteria as cell doublets, thus we designated these cells as secretory progenitors (Sec Pro) (Figure 5A, Figure S7A, B).

**Figure 5.**
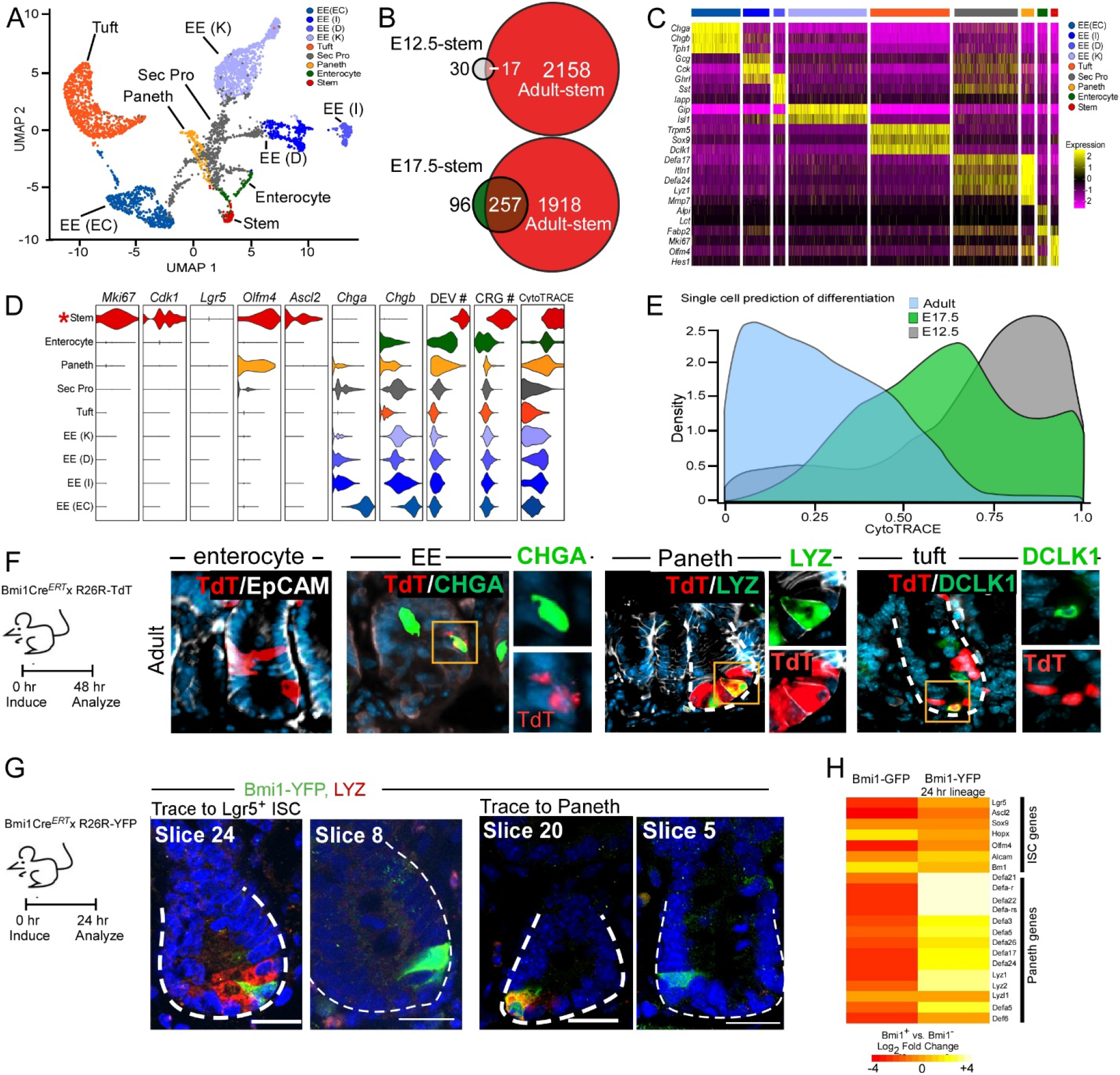
Adult homeostatic intestines harbor Bmi1^+^ ISCs. (A) UMAP plot of adult Bmi1-GFP^+^ cells annotated by their intestinal cell type. (B) Differential expressed genes between E12.5 or E17.5 Bmi1^+^ ISC population versus adult ISC population, logFC > 2, FDR < 0.01. (C) Heatmap of differentially expressed marker genes for intestinal subpopulations among Bmi1-expressing cells. (D) Violin plots of intestinal stem cells (ISC, *Mki67, Cdk1, Lgr5, Olfm4, Ascl2*), and enteroendocrine cells (EE, *ChgA, ChgB*) related genes and developmental score (DEV #), chromatin regulatory gene score (CRG #), and CytoTRACE score. (E) Distribution of CytoTRACE scores for E12.5, E17.5, and Adult Bmi1-GFP^+^ cells. (F) Short term, 48 hr TdT-lineage tracing from the *Bmi1* promoter in adult mice give rise to enterocytes (expressing EpCAM), EE cells (expressing chromograninA, CHGA), Paneth precursors (expressing lysozyme, LYZ) and tuft cells (expressing doublecortin like kinase 1, DCLK1). (G) Intestinal crypts analyzed for lineage in short term, 24 hr, YFP-lineage tracing from the Bmi1 promoter in adult mice. Confocal analyses of YFP and stained lysozyme (LYZ) expression in two crypts. (Left) Two YFP^+^ cells, with one situated between paneth cells, and (right crypt) Two YFP^+^ cells, with one co-expressing LYZ. (H) FACS-isolated YFP^+^ lineage tracing in (G), cells subjected to qRT-PCR analyses for stem cell and Paneth cell genes.

These observations, along with published reports of pre-terminal EE cells (or secretory precursor cells, Sec Pre) dedifferentiating towards Lgr5^+^ ISC in response to injury^11,13^, support the concept that adult Bmi1^+^ ISCs and Bmi1^+^ epithelial cells in general may retain developmental plasticity. In this scenario, Bmi1^+^ cells may be capable of executing an earlier developmental program to re-engage proliferative stem cell phenotypes in response to injury or disease states. Therefore, to explore this idea, we performed DEG analyses between E12.5 and adult Bmi1^+^ scRNA-seq datasets (Table S5). We designated the top 50 DEGs that were upregulated in E12.5 versus adult Bmi1^+^ cells as a developmental score (DEV). To expand analyses of stemness, we also established a score for genes involved in chromatin regulation using the same approach (Table S6), calling this measure the chromatin regulatory gene (CRG) score^59,60^. We then evaluated each adult Bmi1^+^ cell cluster for hallmarks of stemness including discrete stem cell gene expression (*Mki67, Cdk1, Lgr5, Olfm4, Ascl2*), DEV score and CRG score, and found only the adult Bmi1^+^ stem cell population had discernable stem cell identity (Figure 5D). Importantly, the adult stem cell population did not express significant levels of *Chga* or *Chgb*, further distinguishing it from mature EE cells, preterminal EE cells, EE progenitors or secretory progenitors^11,13,44^.

A published scRNA-seq study examined different adult ISC populations using GFP reporter mouse models, including Bmi1-GFP^+^, Prox1-GFP^+^, Lgr5-GFP^+^, and Lgr5-GFP^-^ cells^16^. The combined analyses demonstrated that the majority of Bmi1-GFP^+^clusters separated from other cell populations and expressed a variety of EE cell genes. However, a small number of Bmi1-GFP^+^ cells were positioned away from EE gene-expressing cell clusters and lacked differential EE protein expression^16^. Our independent re-analyses of this scRNA-seq Bmi1-GFP^+^ dataset (Yan et al dataset) confirmed that these previously incongruous Bmi1^+^ cells form their own cluster (Figure S6B), similar to what is found in our dataset (Figure 5A). This cluster possessed similar gene expression, DEV, and CRG scores to the Bmi1^+^ stem cluster in our adult Bmi1^+^scRNA-seq dataset (Figure S5D). After integrating the previously published dataset^16^ with our own (Smith et al dataset), and performing dimensionality reduction analysis, we found that our Bmi1^+^ ISC cluster was superimposed upon the published Bmi1^+^ ISC cluster (Figure S6C), further supporting the similarity of our identified Bmi1^+^ ISC cluster to the published Bmi1^+^ ISC cluster.

To further support the stem identity, we used CytoTRACE to measure the developmental potential of Bmi1^+^ cells. When applied to E12.5, E17.5, and adult Bmi1 ^+^cells, CytoTRACE results aligned each time point as anticipated, with a higher CytoTRACE score predicting a less differentiated state (i.e. E12.5>E17.5>adult, Figure 5E). When we separated CytoTRACE scores by adult Bmi1^+^ cell types, we found that the stem population had the highest score, signifying a greater developmental potential (Figure 5D, far right column).

To confirm stem cell attributes and determine if adult Bmi1^+^ ISCs could give rise to differentiated progeny or to Lgr5^+^ ISC, similar to development, we conducted short-term lineage tracing in intestines from Bmi1Cre^ERT^ x R26R-TdT (or Bmi1Cre^ERT^ x R26R-YFP) mice. We examined lineage-induced intestines after 24 and 48 hrs for co-expression of the Bmi1-derived lineage marker, TdT and differentiated lineage proteins for enterocytes (EpCAM), EE (CHGA), Paneth (LYZ) and Tuft (DCLK1) cells and found that Bmi1-expressing cells traced to all of these differentiated lineages (Figure 5G). To refine the lineage relationship, we used a different experimental paradigm to demonstrate that lineage^+^ (YFP) cells reside in crypts with multiple lineage positive epithelial cells. We performed confocal microscopy of entire crypts 24 hr post-lineage induction, where only 2-3 lineage^+^ cells exist (Figure 5G). In these crypts, we identified co-positive cells juxtaposed to a Bmi-YFP^+^, lineage negative cell, indicating that the lineage^+^ cell was likely derived from the Bmi1-YFP^+^ cell. To support this finding, we then FACS-isolated Bmi1-YFP 24 hr-lineage cells and subjected them to qRT-PCR for Lgr5^+^ISCs and Paneth cell gene expression profiling (Figure 5H). When compared to isolated Bmi1-GFP cells, we observed an increase in gene expression for both Lgr5 ISC^61^ and Paneth cells^62^, demonstrating that adult Bmi1^+^ ISC can function like their developmental counterpart and give rise to both differentiated lineages and Lgr5^+^ ISCs. Collectively these data indicate that Bmi1^+^ ISCs establish multiple lineages that retain *Bmi1* gene expression, across development and in the mature intestine (i.e., E12.5 ISCs lineage trace to EE cells seen in Figure 3E and Figure S5C; E17.5 ISCs lineage trace to enterocytes, Paneth, and EE cells seen in Figure 3E, Figure S5E; and adult ISCs lineage trace to enterocytes, Paneth, tuft, EE cells as seen in Figure 5F, G).

In summary, a multi-faceted analyses of Bmi1^+^ cells across development and adulthood with single cell resolution identify a discrete subset of adult homeostatic intestinal Bmi1^+^ ISCs, that was previously overlooked in bulk –omics analyses of the Bmi1^+^ population^11^. Although slow-cycling^23^, these cells maintain undifferentiated, developmental-like features and gene expression preserved from analogous E12.5 Bmi1^+^ ISC populations (i.e. DEV score) indicating that they could be poised to revert to a more developmental state in the face of challenge.

### In the adult intestine, irradiation induces the emergence of Bmi1-expressing cells with enhanced developmental expression

Despite the radiation-induced elimination of Lgr5^+^ ISCs, the intestine quickly replenishes its pool of progenitor cells while maintaining barrier integrity. The exact identity of the population(s) and mechanisms responsible for this resiliency remains somewhat muddled, as recent studies describe multiple different cell types or underlying mechanisms responsible for post-injury regeneration, including reserve stem populations^20,50^, differentiated or lineage committed cells^11,20,51^ contributing to the plasticity, revival stem cells^63^, and adaptive differentiation^64^, while unfortunately dismiss coordinated mechanisms. To explore a role for Bmi1^+^ ISCs in regeneration after irradiation injury, we and others^12,16^ found a modest expansion in Bmi1^+^ cells in the intestine (Figure 6A). qRT-PCR analyses of homeostatic and irradiated Bmi1-GFP^+^ cells displayed altered expression of stem (e.g. *Lgr5, Bmi1, Hopx*) and differentiated (e.g., *Chga, Chgb*) cell gene expression, as well as a robust increase in expression of developmental genes (e.g., *Trop2, Cnx43, Spp1;* Figure 6B). We confirmed this increase in developmental gene expression by antibody staining for the developmental protein TROP2, which overlapped Bmi1-marked lineage only in irradiated tissue (Figure 6C).

**Figure 6.**
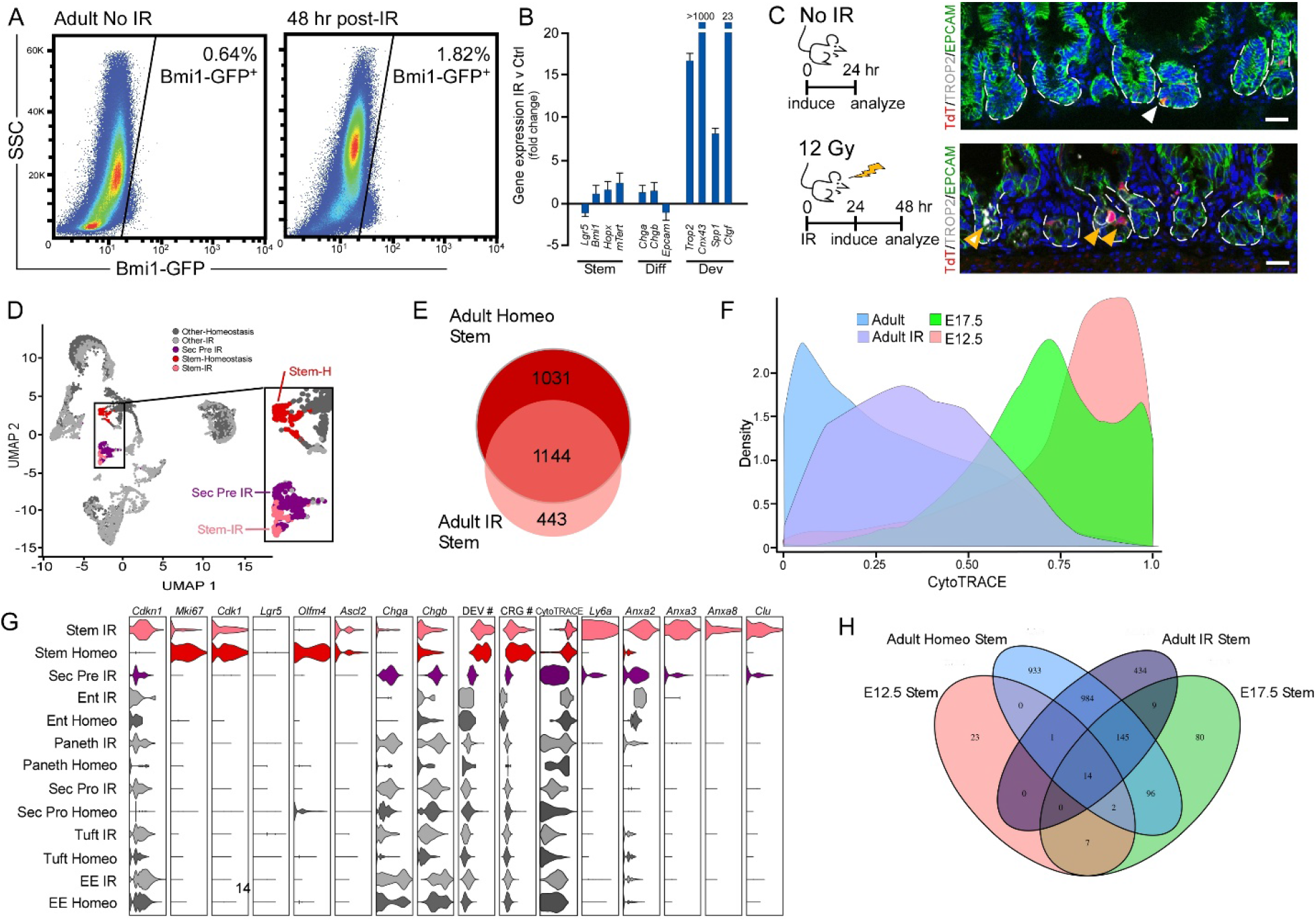
Bmi1^+^ cells express developmental markers in response to injury. (A) FACS plots of adult Bmi1-GFP^+^intestinal epithelial cells at homeostasis (left) or 48 hr after 12Gy radiation (IR, right). (B) qRT-PCR analysis of gene expression in Bmi1-GFP^+^ FACS-isolated cells 48 hr post-IR relative to non-irradiated controls. Representative data from technical replicates. Gene expression analyses were performed on N=4 independent experiments representing at least N=8 mice. Data are presented as mean ± SEM of triplicate fold change. (C) Small intestinal sections from Bmi1-Cre TdTomato (TdT) un-IR control mice (left) and 48 hr post IR (right) stained with antibodies against EPCAM (green) and TROP2 (white), with TdT expression in red. White arrowheads denotes Bmi1-TdT cells, yellow arrowhead denote TdT cells co-expressing TROP2. White dashed lines denote the epithelial-mesenchymal boundary. Scale bar = 25 μm. N=4 total mice per group. (D) UMAP visualization of Bmi1-GFP^+^ cells from adult un-IR and adult 48 hr post IR intestines annotated by cell type. Un-IR and IR Bmi1-expressing subpopulations are largely aligned. One cluster was exclusively composed of irradiated cells (boxed) and contained cells with differential stem and secretory gene expression. (E) Differential expressed genes between adult Bmi1^+^ Stem-homeostasis versus Stem-Irradiated populations, logFC > 2, FDR < 0.01. (F) Distribution of CytoTRACE scores for E12.5, E17.5, adult, and adult IR Bmi1-expressing cells. (G) Violin plots of DNA damage (*Cdkn1*), intestinal stem cell (ISC; *Mki67, Cdk1, Lgr5, Olfm4, Ascl2*), enteroendocrine cell (EE; *Chga, Chgb*) gene expression, developmental (DEV #), chromatin regulatory gene (CRG #), and CytoTRACE scores reveal similarity between un-IR (pink) and IR (red) stem populations and dissimilarity with IR secretory precursor (purple) population. The IR stem population uniquely expressed developmental genes (*Ly6a, Anxa2, Anxa3, Anxa8, Clu*). (H) Venn diagram showing overlapping of DEGs of identified stem cell clusters in E12.5, E17.5, adult homeo and adult IR Bmi1-GFP^+^ datasets.

To discern which Bmi1^+^ subpopulations might contribute to intestinal repair of the intestine, we performed scRNA-seq on adult Bmi1-GFP^+^ cells that were FACS-isolated 48 hours after irradiation (IR). We combined a scRNA-seq dataset of 3,607 Bmi1^+^ IR cells with our homeostatic adult Bmi1^+^ scRNA-seq dataset to identify transcriptional differences between homeostatic and irradiated Bmi1^+^ cells. We employed dimensionality reduction, clustering, and differential gene expression analysis to annotate the cell types (Table S8). Analogous cell identities detected in the homeostatic and irradiated datasets were superimposed in UMAP visualization (Figure 6D). However, the homeostatic stem population (Stem Homeo) did not overlap with the irradiated stem clusters that was composed of two discrete subpopulations (i.e., Stem irradiated: Stem-IR, and the pre-terminal EE population, secretory precursor: Sec Pre IR). Comparison of DEGs revealed that the Bmi1^+^ Stem Homeo population shared 53% of expressed genes with the Stem IR subpopulation (Figure 6E). Further, the Stem IR subpopulation lacked gene expression patterns observed in the homeostatic ISC population, specifically *Ki-67, Cdk1*, and *Olfm4* (Figure 6G), but retained a similar expression of the Wnt target gene, *Ascl2*. In addition, the Stem IR cluster had moderate expression of *Cdkn1, a* gene that encodes p21, a cyclin dependent kinase inhibitor, that when expressed at low levels allows cells to proliferate^65^ (Figure 6G). Furthermore, key developmental stem cell genes were differentially expressed in the Bmi1^+^ Stem IR population, that we identified in E12.5 Bmi1^+^ subpopulations, including *Anxa1-5, Anxa11*, and *Clu* (Table S2, Figure 6G). This indicates that the Bmi1^+^ stem population undergoes discrete alterations in response to radiation injury and in the absence of the Lgr5^+^ ISC population. The Sec Pre IR subpopulation clustered closely with the Stem IR subpopulation, and harbored EE-related gene expression (including *Chga, Cck, Ghrl, Nts, NeuroD1* and *Serapina 1c*; Figure 6G, Figure S7C). This subpopulation was similar to the reported Bmi1-expressing <u>sec</u>retory <u>pre</u>cursor population^11^ described as a pre-terminal EE or EE precursor capable of dedifferentiation into an Lgr5^+^ ISC in response to injury.

To further assess the developmental identity of the Stem IR and Sec Pre IR subpopulations, we evaluated them for CytoTRACE, DEV and CRG scores (Figure 6F, G), and found that the Stem IR and Stem Homeo subpopulations were nearly identical, while the Sec Pre IR population had lower scores across these three metrics. Further, Stem IR and Sec Pre IR populations harbored unique gene expression when compared side by side, including YAP target genes *Ly6a, Anxa8 and Clu* that repress the canonical Wnt pathway (Figure 6G, Figure S7C). We could interpret these differences to be that the Stem IR population expresses developmental signatures by repressing the canonical Wnt signaling pathway. Further, it is possible that the Sec Pre IR population functions by an independent mechanism. While these two populations have distinct expression profiles and predicted developmental potential their close proximity indicates related identity in response to radiation injury. Despite this, only the Stem IR population had analogous Stem Homeo features.

To directly identify differences between Bmi1^+^ISC populations at different states (i.e., development, homeostasis and in response to irradiation), we compared DEGs from cluster 0 (E12.5, light gray), cluster 0,4 (E17.5, dark green), adult homeo stem and adult IR stem subpopulations (Figure 6H, Table S3). Both adult Bmi1^+^ ISC subpopulations shared more expressed genes with the E17.5 than the E12.5 ISC population, with a majority of these genes representing functions involving regulation of the proliferation. Surprisingly, despite an enhanced developmental gene expression pattern, the adult stem IR subpopulation did not have greater overlap with E12.5 DEGs. Interestingly, all stem populations share a core of 14 expressed genes that represent regulation of the mitotic spindle, epigenetic regulation, or protein degradation. These comparisons indicate that adult Bmi1^+^ ISC populations share a level of identity with their developmental counterparts. Further, emergence of a developmental profile superimposed onto the gene profile of the homeostatic Bmi1^+^ ISC indicates that re-expression of a proliferative developmental state within the Bmi1^+^ ISC may facilitate or participate in tissue regeneration in this context. Reported reserve stem populations^15,66,67^, revival stem cells^63^, and de-differentiating cell types^11,13,20^ are implicated in driving epithelial recovery in response to injury; our data identifies a third mechanism for tissue regeneration—specifically harnessing a latent developmental state.

### Bmi1-expressing epithelia in tumor environments co-express developmental markers

Bmi1 is a prognostic marker for colorectal cancer, with higher expression being associated with decreased survival^68,69^, as well as a marker of colorectal cancer stem cells^70^. Given the stem identity of a subpopulation of Bmi1-expressing cells in the adult intestine, we probed murine tumors from Apc^Min^ animals for Bmi1, Trop2, and β-catenin protein expression and found that Bmi1^+^ tumor cells co-expressed Trop2 but lacked β-catenin expression (Figure 7A). Further, isolated Bmi1^+^ cells from Apc^Min^ tumors grew as spheroids (Figure 7B), reminiscent of the spheroids grown from the canonical Wnt-indepdent^23^, developmental intestine (Figure 1A). This finding indicated that Bmi1-expressing tumor cells harbored developmental identity, but with canonical Wnt signaling independence.

**Figure 7.**
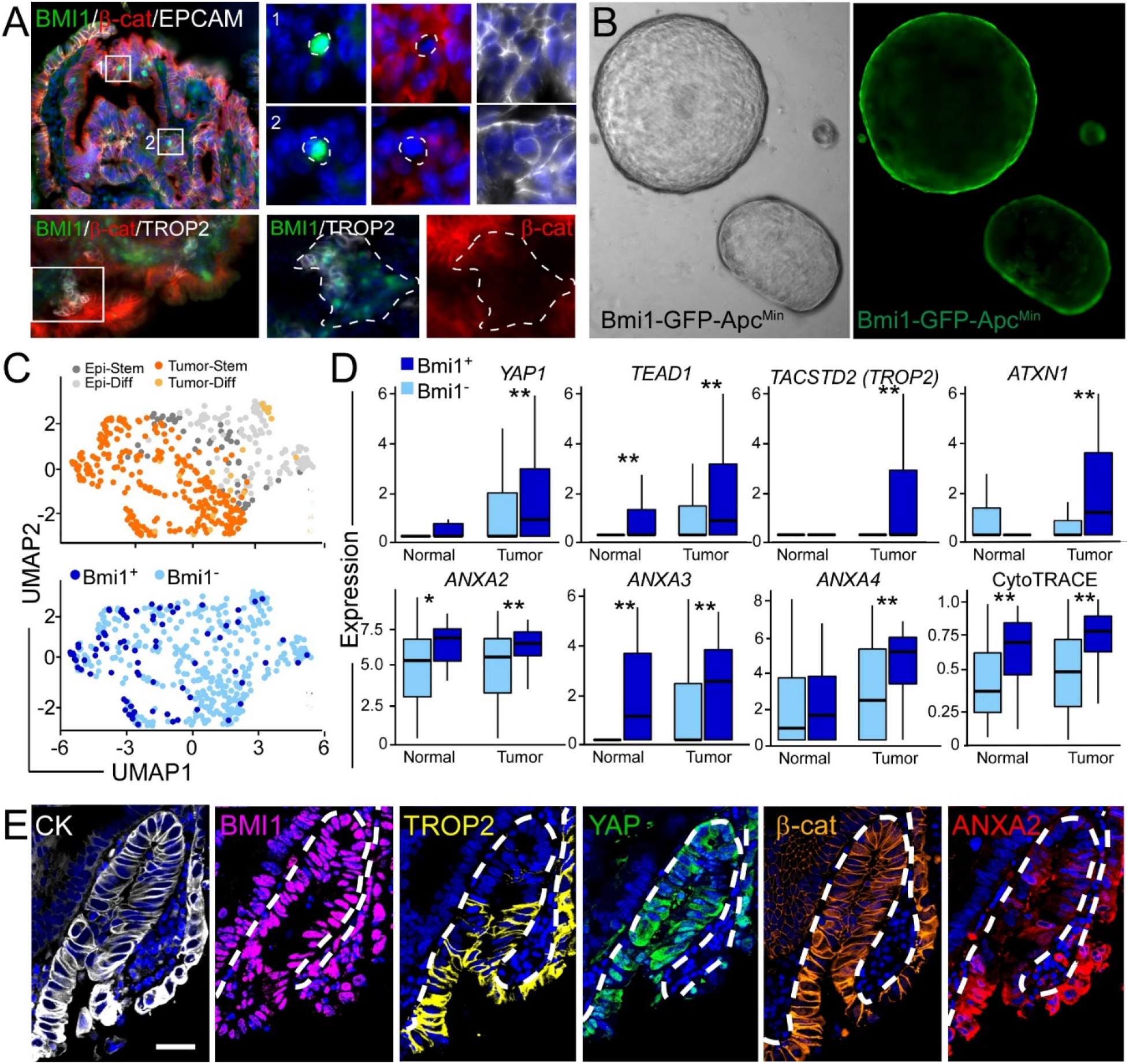
Bmi1^+^ tumor cells maintain developmental-like expression. (A) IHC images of murine intestinal adenomas reveal resident BMI1/β-catenin/EPCAM and BMI1/TROP2/EPCAM co-expressing cells. (B) Developmental-like spheroids derived from Bmi1-GFP-Apc^Min^ tumor cells. (C) UMAP visualization of normal epithelial (Epi) and colorectal cancer (Tumor) cell scRNA-seq data annotated by their published cell type (top) and expression of Bmi1 (bottom). Majority of Bmi1-expressing cells reside in normal and tumor stem populations. (C) Boxplots reveal higher developmental gene (*YAP1, TEAD1, TACSTD2, ATXN1, ANXA2, ANXA3, ANXA4*) expression and CytoTRACE scores of Bmi1-expressing cells in both normal and tumor tissue. (* P-value < 0.02, ** P-value < 0.003) (D) Human intestinal adenocarcinoma images highlight Bmi1-expressing cells that co-express developmental proteins (TROP2, YAP, β-Cat, ANXA2).

To determine if this paradigm extended to human disease, we profiled an annotated human scRNA-seq dataset with cells from 11 primary colorectal cancer tumors and adjacent normal mucosa^71^. Focusing on tumor and adjacent normal epithelial cells (272 tumor cells and 160 normal cells) we categorized cells as stem/cancer stem or differentiated, then analyzed the entire population for *BMI1* RNA expression, with the majority of tumor cells possessing stem cell identity (Figure 7C). Cells with detectable *BMI1* RNA were predominantly in the stem cell clusters. To determine if *BMI1^+^* tumor cells differentially harbored developmental profiles we compared developmental gene expression and applied CytoTRACE between *BMI1*-expressing and non-expressing cells (Figure 7D). *BMI1^+^* cells in both adjacent normal and tumor tissue consistently had higher undifferentiated cell identity as shown for CytoTrace score (p-value < 0.002). Validating this observation, Bmi1-expressing cells in human colorectal cancer tissue also expressed significantly higher developmental protein expression (e.g., Trop2, Anxa2, and Yap, p-values < 0.05) (Figure 7D). Finally, we observed increased expression of developmental proteins in human CRC tumors (Figur 7E). These findings indicate Bmi1-expressing tumor epithelia harbor developmental identity analogous to expression in highly proliferative embryonic and injury stimulated Bmi1^+^ ISCs.

## Discussion

Slow-cycling, label retaining cells within the intestinal crypts historically were described as ISC^72^ until the discovery of the more rapidly proliferating, canonical Wnt-responsive Lgr5^+^ ISC population^1^. Since this discovery, the intestinal stem cell field has identified myriad different stem cell populations and mechanisms for safe guarding epithelial integrity during development, homeostasis, and regeneration—and thus historically defined ISC have come under intense scrutiny. The once-recognized, slow-cycling Bmi1^+^ ISC is one such population. Bmi1 epithelial expression locates to rare crypt-base cells in slow-cycling position, but is also detected in differentiated EE cells in the adult intestine during homeostasis^12,16^. Further, a recent study found that Bmi1^+^ EE precursor cells have plasticity to dedifferentiate toward Lgr5^+^ ISCs^11,13^ and accompanying bulk Bmi1^+^ population suggested that they lack stem cell properties^22^, thus calling into question the authenticity of stem cell properties within Bmi1-expression epithelia.

Work from our group has long supported a stem cell role for the Bmi1^+^ epithelial population^23^, and thus we set out to more deeply examine the population. We evaluated stem cell states in the Bmi1^+^ epithelial population across intestinal development, adult homeostasis, in response to injury, and in tumorigenesis. Using a combination of murine models, lineage tracing, human tissue and -omics analyses of Bmi1^+^ populations with single cell granularity, we established that discrete subpopulations harbor stem cell attributes. Across development of the intestine, two discrete stem cell states of Bmi1^+^ cells exist, the first in early development characterized by robust proliferation, expression of pioneer transcription factors and hox genes, and the second stem cell state, in late embryonic development, as a minor population of slow-cycling cells fundamentally different from its earlier developmental counterpart. Similar to late intestinal development, adult tissue at homeostasis harbors a minor population of Bmi1^+^ cells with expression of non-canonical Wnt signaling target genes, and a high CytoTRACE score aligning it with developmental identity. Moreover, when isolated and grown in culture, adult tissue Bmi1^+^ cells recapitulate the *in vitro* phenotype of developmental Bmi1^+^ ISCs^23^. Finally, in response to injury we identified two discrete subsets of Bmi1^+^ cells with stem features, the first (stem IR) aligned with the homeostatic Bmi1^+^ ISC population, except that it differentially expressed high levels of developmental genes (i.e. *Clu, Ly6a*), as well as previously published “reserve” cell gene expression^63^. The second discrete subpopulation (Sec Pre) expressed a gene signature aligned with the previously reported secretory precursor population identified by bulk RNA-seq analyses^11^, that is thought to be the product of dedifferentiated EE cells, which ultimately contributed to regeneration of the active cycling, canonical Wnt-dependent Lgr5^+^ ISC pool. Whether these cells are dedifferentiated from Bmi1^+^ EE cells, or are differentiating population emerging from the stem IR population cannot be determined by our data. Emergence of Bmi1^+^ ISCs with early developmental gene expression profiles may indicate that even during homeostasis, Bmi1^+^ ISCs are poised to re-engage developmental identity to re-establish the active-cycling stem cell niche in response to injury, bringing forth an additional mechanism, rooted in development, for tissue regeneration.

During early intestinal development, the proliferative Bmi1^+^ ISC population exists in high numbers. It is the predominant stem cell population at E12.5, at a time when canonical Wnt-dependent, Lgr5^+^ ISCs are not yet present. The temporal relationship with Lgr5^+^ ISCs along with lineage tracing studies reveal that Bmi1^+^ cells trace to Lgr5^+^ISCs. Additionally, short-term lineage trace from the Bmi1^+^ promoter, initiated at E12.5 or E17.5 revealed relationships between the Bmi1^+^ ISC and multiple epithelial lineages (i.e. Paneth, EE and enterocyte), indicating that Bmi1^+^ ISCs are capable of contributing to downstream lineage independent of Lgr5^+^ cells. scRNA-seq analyses of E12.5 and E17.5 FACS-isolated Bmi1^+^ epithelia identified secretory and enterocyte gene expression, thus corroborated lineage-tracing findings. Interestingly, these lineage relationships with Bmi1^+^ ISCs were also identified in adult homeostatic tissue, with the observation of short-term lineage tracing to enterocytes, Paneth, EE and tuft cells. Moreover, Lgr5^+^ ISCs were also among Bmi1^+^ progeny. This finding indicates that adult homeostatic Bmi1^+^ ISCs also harbor the ability to generate Lgr5^+^ ISCs during homeostasis to replenish the Lgr5^+^ ISC pool. Our finding expands on the identification of a “revival” stem cell that expresses developmental genes in response to injury and can give rise to Lgr5^+^ cells^63^, adding evidence that a similar Bmi1^+^ ISC, without developmental gene expression, can similarly function during homeostasis, but its developmental predecessor can contribute to repair in response to injury. Interestingly, our finding offers an unappreciated mechanism for a background replenishment of the active-cycling stem cell pool and low level of lineage differentiation during homeostasis.

We identified a potential role for non-canonical Wnt signaling in regulating a developmental Bmi1^+^ ISC population. Non-canonical Wnt signatures are highly indicated in scRNA-seq and ATAC-seq datasets from early developmental time points. These signatures wane as the Bmi1^+^ ISC population become less proliferative. This coincides with the expression of canonical Wnt ligands in the developing intestine^31,73^ and establishment of Lgr5^+^ ISC. The canonical Wnt signaling microenvironment correlates with the developmental Bmi1^+^ ISC becoming less more quiescent and is similar, but perhaps transitional, to the adult Bmi1^+^ ISC population, as E17.5 Bmi1^+^ ISC cells share 72% of their genes with the adult Bmi1^+^ ISC population. However, although it is intriguing to hypothesize differential impact from canonical versus non-canonical Wnt signaling on Lgr5^+^ and developmental Bmi1^+^ ISC populations, the discrete regulatory control of Bmi1^+^ ISC’s proliferative transition remains unknown. Insights into how stem cell populations re-engage developmental memory and gene expression will open the potential ability to control the impact of developmental gene expression in cancer and in tissue regeneration.

In summary, we show that discrete subpopulations of Bmi1^+^ cells contain bona fide stem cell attributes across intestinal development, into adulthood and in response to injury or disease. Moreover, we provide evidence that a non-canonical Wnt signaling pathway may regulate early developmental Bmi1^+^ ISC proliferation, setting up a regulatory dichotomy between Wnt-responsive Lgr5+and Bmi1^+^ ISC populations. We present evidence that Bmi1^+^ ISCs can lineage trace to Lgr5^+^ ISCs during development, but importantly also in the homeostatic intestine. This indicates that the Bmi1^+^ ISC population serves as a resource for replenishing the Lgr5^+^ ISC population during homeostasis. Further, Bmi1^+^ ISCs could contribute to the myriad mechanisms that protect epithelial integrity in response to injury or are dysregulated in diseases such as cancer. Our finding establishes an underappreciated mechanism for Bmi1^+^ ISC contribution to intestinal development and homeostasis, and therefore highlights the complicated but robust interwoven network of diverse stem, progenitor and cellular plasticity that provides redundant mechanisms to safeguard the integrity of organs—like the intestine—essential for life.

## Acknowledgements

We are grateful for support from the Advanced Light Microscopy Core (ALMC) and Stefanie Petri-Kaech, director of the ALMC. We thank Austin Nguyen, Benjamin Weeder, and Abhinav Nellore for their insightful comments and Thomas Sutton for guidance on statistical analyses. This work was supported by CA069533, DK085525, DK068326.

## Methods

### Mouse strains and animal studies

All experiments using animals were approved by the OHSU IACUC committee. Mice were housed in specific pathogen-free barrier within the OHSU vivarium under a continuous 12 h day/night cycle with access to animal chow (5001, PMI Nutrition International, Richmond, IN) and water ad libitum. Adult mouse experiments were conducted with animals between 8-12 weeks of age with equal contribution from each gender and used litter-mate controls when appropriate. For radiation-induced injury, a cesium irradiator was used to deliver a 12Gy whole-body dose. Lineage tracing in intestines from adult mice or embryos was initiated with delivery of Tamoxifen (#T5648, Sigma, St. Louis, MO) dissolved in corn oil at a dose of 0.08 mg/g body weight by intraperitoneal (IP) injection or oral gavage to pregnant dams, respectively. The following mouse strains were obtained from the Jackson Laboratory (Bar Harbor, ME): C57Bl6/J (#000664), Bmi1-GFP (#017351)^7^, Bmi1-CreERT2 (#010531)^74^, Lgr5-GFP-IRES-CreERT2 (#008875)^1^, ApcMin (#002020)^75^, and R26R-Tomato (#007909).^76^

For developmental studies, timed mating pairs were established and pregnant dams sacrificed to dissect embryonic intestines at embryonic day (E)12.5-19.5. Bmi1- and Lgr5-GFP embryos were screened by wholemount fluorescence of GFP within the developing brain or otic cup, respectively, on a Leica DMIRB inverted fluorescent microscope. Embryos fostered to adulthood were isolated at E19.5 and fostered into C57Bl6/J litters.

### Statistics

Data are presented as mean ± standard error of the mean (SEM). Statistical significance was determined using an un-paired student’s t-test with Welch’s correction. A p-value < 0.05 was deemed statistically significant. Statistical analyses were performed using Prism software (GraphPad, La Jolla, CA). All experiments were performed with n = 4-12 total mice with at least two independent technical experimental replicates.

### Tissue isolation and preparation

Adult intestinal tissues were isolated and prepared as previously described.^77^ Briefly, adult mice were euthanized, intestinal tissues rapidly isolated, luminal contents flushed with ice-cold PBS and tissues fixed by incubation with 4% PFA in PBS for 1hr at room temperature or further prepped for subsequent epithelial cell dissociation for Flow cytometry or FACS.^23^ Developmental tissue samples were isolated and prepared similar to previous reports.^78^ Dams were euthanized and embryos quickly isolated and genotyped as described above. For E12.5 embryos, the whole embryo was fixed in 2% EM-grade PFA in PBS (Electron Microscopy Sciences) for 1hr at room temperature or intestines were isolated by dissection under wholemount microscope (Leica microsystems) for dissociation and subsequent flow cytometry or FACS. For E14.5 and later samples, the small intestine was isolated by dissection under wholemount microscope and fixed in 2% EM-grade PFA as above or dissociated.

### Immunohistochemistry and microscopy

Fixed tissue samples were embedded in Tissue Tek OCT compound (Sakura Finetek USA), and 5-10um sections were collected onto glass slides in a cryostat (Leica biosystems). Tissue sections were washed in PBS, blocked using PBS-blocking buffer (PBS containing 1% BSA, 1mM CaCl2, and 0.1% Triton X-100), and subsequently stained with antibodies (Table S1). Slides were mounted in n-propyl gallate solution and imaged on a Zeiss ObserverZ1 microscope with ApoTome optical sectioning (Carl Zeiss), and digitally scanned on a Zeiss Axio Scan.Z1. The filter cubes used for digital image collection were DAPI (96 HE BFP), AF488 (38 HE Green Fluorescent Prot), AF555 (43 HE DsRed), AF647 (50 Cy 5). The exposure time, light source intensity for the digitally scanned, fluorescently labeled lineage traced slides was as follows: 20 ms, 20.00% (DAPI), 10 ms, 50.00% (AF488), 350 ms, 75.00% (AF555), and 100 ms, 75.00% (AF647). Full tissue scans were taken with the 20x objective (Plan-Apochromat 0.8 M27, Zeiss). Image analyses were performed using ImageJ (National Institutes of Health) and Zeiss Zen blue software (Carl Zeiss AG, Germany).

### Flow cytometry/FACS

Adult intestinal epithelium was isolated, crypt-enriched and dissociated to a single cell suspension as described previously.^23^ Developmental samples were dissociated by mincing whole small intestinal tissues in 1x TrypLE-express digestion reagent (Gibco) supplemented with 50ug/mL DNAseI (Roche) and 10 μM Y-27632 (Stemgent) and incubated at 37 °C for 5-10 min with occasional pipetting with a P-1000 pipette tip, followed by filtration through a 35 μm cell-strainer lid (BD Biosciences). Resultant cell suspensions were stained with antibodies (Table S1) and sorted for purity on a BD Influx cell sorter at the OHSU Flow Cytometry Shared Resource. Flow cytometry and FACS analyses were performed using FlowJo Software (TreeStar).

### *Ex vivo* cell culture

3D *ex vivo* cultures were established using previously described methods^24^. Briefly, cell populations were isolated and embedded in Matrigel containing the following recombinant growth factors: Egf (Peprotech), Noggin (Peprotech), R-Spondin1 (R&D Systems). The solidified Matrigel-cell mixture was then overlaid with base culture media: Advanced DMEM/F12 containing 10mM HEPES (Gibco), N2 and B27 nutrient supplements (Gibco), Pen/Strep (Gibco) and 10uM Y-27632 (Stemgent).

### qRT-PCR

RNA from FACS-isolated cell populations or cell cultures were isolated using the RNeasy mini kit (Qiagen) with on-column DNAse digestion according to the manufacturer’s protocol. 1ug of purified RNA was converted into cDNA using the High-capacity cDNA synthesis kit (Applied Biosystems) and the quality assessed on a Bio-analyzer (Agilent). qRT-PCR was performed on a Viia7 (Applied Biosystems) using Taq-Fast (Applied Biosystems) or Sybr green (Qiagen) reagents and the probes/primer sets listed in Table S2. Gene expression was quantified by the ΔΔCt method^79^ using GAPDH as the internal reference gene. Results are mean ± SEM from triplicate analyses. Heatmaps were generated using the Complex Heatmap package in R with the default parameters.^80^

### ATAC-Seq

For bulk ATAC-Seq, intestinal epithelial cell populations were FACS-isolated directly into FACS buffer (DMEM/F12, 0.5% dialyzed FBS, 1% HEPES, 0.02% Y-27632 (Sigma-Aldrich, MO) from N=6 embryos at each time point. Cells were then isolated using 1 mL ice cold Nuclei Isolation Buffer (NIB; 10 mM HEPES-KOH, pH 7.2 [Fisher, Cat. BP310-1], 10 mM NaCl [Fisher, Cat. M-11624], 3 mM MgCl2 [Sigma, Cat. M8226], 0.1% IGEPAL [v/v; Sigma, I8896], 0.1% Tween-20 [v/v, Sigma, Cat. P7949], and 1x protease inhibitor [Roche, Cat. 11873580001]) with trituration followed by pelleting at 500xg, removal of the supernatant, and resuspension in 200 μL NIB. Nuclei were then quantified using a hemocytometer and then diluted to 10,000 nuclei per 1 μL. 3 μL of nuclei (30,000) were added to 14.5 μL of NIB with 2.5 μL of 4X TAPS tagmentation buffer (4X TAPS-TD buffer (132 mM TAPS pH=8.5, 264 mM potassium acetate, 40 mM magnesium acetate, and 64% dimethylformamide). 5 μL of 500 nmol loaded Tn5 transposase with standard Nextera adapters was then added and incubated at 55 °C for 5 minutes, then placed on ice prior to reaction stopping and cleanup using a QIAGEN MinElute PCR Cleanup column workflow, eluting in 10 μL EB. 25 μL of Kapa HiFi Non hot start PCR master mix was then added with 2.5 μL forward and reverse primers at 10 μM, with the reverse primer containing a 10 bp index, 0.5 μL 100X SYBR Green and 12 μL water. PCR was then performed using the following: 72 °C for 10:00, 99x(0:30 at 98 °C; 0:30 at 65 °C; 1:00 at 72°C; plate read) on a BioRad CFX real time thermocycler. Reactions were stopped when exponential amplification was observed at 12 cycles. Libraries were then cleaned up using AMPure beads, quantified on an Agilent Bioanalyzer, and sequenced on an Illumina NextSeq 500.

Sequence reads were demultiplexed, allowing for 1 hamming distance in the library index followed by alignment to the reference genome GRCh38 using bwa mem. Reads were duplicate-removed using samtools rmdup and then peaks called using macs2 using default parameters. Peaks were called for each individual dataset as well as aggregated across all data and the union peak set was used for all subsequent analysis. A count matrix was then produced of ATAC reads for each peak for each sample and DESeq2 was then used to identify differentially accessible peaks, using the comparison sample sets detailed in Fig. 4H. Motif enrichment was carried out using ChromVAR to produce global motif deviation z-scores for each transcription factor motif. Wnt-related factors (i.e. known ligands or inhibitors^73^) were then selected for plotting to reveal trends across time points and clustered using hclust in R.

### Single-cell RNA-sequencing

Developmental and adult epithelial cells were FACS-isolated and used for downstream processing. Single cell RNA-sequencing was performed using the Chromium Single Cell 3’ GEM, Library, and Gel Bead Kit (10X Genomics) and the Chromium Single Cell B Chip Kit (10X Genomics) according to the 10X Chromium version 3 protocol (CG000183). In short, FACS-isolated cell populations suspended in PBS were loaded into 10X chromium microfluidic chips and single cells were encapsulated into Gel Bead-in-Emulsion by the Chromium controller (10X Genomics). RNA from individual cells were barcoded and libraries prepped according to the 10X chromium version 3 chemistry using the C1000 Touch thermal cycler (Bio-Rad). Quality of prepped cDNA and resultant libraries were assessed with the Agilent 2100 Bioanalyzer. Sequencing was performed on the HiSeq 2500 sequencer (Illumina) by the OHSU Massively-Parallel Sequencing Shared Resource core.

### Analysis of scRNA-seq data

Cellranger 3.1.0 (10X Genomics) was used for base call file processing, sample demultiplexing, unique molecular identifier (UMI) collapsing, and alignment. We aligned our FASTQ files to an mm10 genome that included the eGFP reporter sequence. The Seurat R package (3.1.2)^81^ and Scanpy Python package (1.8.0)^82^ were used for subsequent analysis. We retained cells with greater than 200 genes expressed and a mitochondrial content lower than 20 percent. Following preliminary analysis, cell clusters determined to be contaminating pancreas tissue or mesenchymal cells were removed in the E12.5 and E17.5 data. In total, we obtained 4404, 3595, 4315, and 3607 cells for E12.5, E17.5, Adult Homeostatic, and Adult Irradiated datasets, respectively.

We applied the SCTransform function in Seurat (for two adult datasets) and Scanpy API (for E12.5 and E17.5 data) for normalization, scaling, and variable feature identification. We performed principal component analysis (PCA) for dimension reduction after selecting highly variable genes according to the default setting in Seurat and Scanpy. Selected PCs were leveraged in Uniform Manifold Approximation and Projection (UMAP) and clustered by the Leiden algorithm^83^ as implemented in Seurat and Scanpy. For the E12.5 and E17.5, Adult Homeostatic and Adult Irradiated analyses, 50, 50, 11, and 11 PCs were selected and 1.0, 1.0, 1.2, and 1.8 were used for clustering resolution, respectively. To perform CytoTRACE analyses to predict the differentiation state of cells, we used the CytoTRACE web application for individual datasets by uploading gene expression matrices onto https://cytotrace.stanford.edu as described at the web site. For Figure 5D and 6D, we downloaded the R code via https://cytotrace.stanford.edu/CytoTRACE-master.zip and performed the CytoTRACE analysis locally after modifying the code to assign NA to 0 in the similarity_matrix of the similarity_matrix_cleaned function in iCytoTRACE.R. To infer developmental trajectories for the E12.5 and the E17.5 data, we applied the partition-based graph abstract (paga) algorithm^84^ in Scanpy to calculate the connectivity of cell clusters and the diffusion pseudotime (dpt) function^84,85^ to conjecture the temporal progression of these clusters.

To integrate the E12.5 and E17.5 datasets, we used the Harmony approach^37^ with default parameters via Scanpy. To calculate the overlap of these two datasets in the merged UMAP (Figure 4A), we used the fuzzy neighborhood network calculated for UMAP. For each cell in the merged dataset, we checked its neighbors and flagged it as an overlapped cell if the neighbors contain at least one cell from another batch. The number of total overlapped cells is 794, out of 7999 cells in the merged dataset.

We analyzed the previously published murine Bmi1-GFP^+^ scRNA-seq dataset using their outlined workflow. To combine both datasets, we followed Seurat’s integration steps by normalizing each dataset individually with SCTransform before running the SelectIntegrationFeatures, PrepSCTIntegration, FindIntegrationAnchors, and IntegrateData functions that provided an integrated expression matrix for PCA, UMAP, and clustering.

For analyses of the human colorectal cancer scRNA-seq data^71^, the original cell annotations were used to remove non-epithelial cells from the raw count data. Downstream analyses proceeded with the standard workflow, using 13 PCs. All analysis scripts in R and Python are hosted at https://github.com/reactome-fi/bmi_isc_analysis. The generated scRNA-seq data were uploaded into GEO (accession number: GSE212798). The original raw counts files were uploaded into SRA (accession number: PRJNA877285).

### Differential gene expression analysis and gene list scoring

Differential gene expression analysis was used to annotate the clusters identified after the scRNA-seq analysis workflow. The Seurat functions FindAllMarkers and FindMarkers were run with default settings for two adult scRNA-seq datasets, and the rank_genes_groups function in Scanpy was used for the E12.5 and E17.5 dataset. Gene list scoring was accomplished with the Seurat function AddModuleScore.^60^ To annotate cell clusters, we used the markers for cell types in Table S3.

### Reactome pathway analysis

We used the annotated pathways in Reactome^54^ (Release 79, December 2021) for pathway activity analysis. The mouse pathway gene set file (https://github.com/reactome-fi/single-cell-analysis/blob/master/resources/MouseReactomePathways_Rel_79_122921.gmt) was generated from mouse pathways, which were computationally predicted based on manually annotated human pathways according to homologous mappings between human genes and mouse genes. To infer individual cells’ pathway activities, we utilized pySCENIC package^86^’s AUCell^53^ function wrapped in Reactome’s scpy4reactome package (https://github.com/reactome-fi/fi_sc_analysis/tree/reactomefiviz_release_8.0.0/python).

## Notes

https://github.com/reactome-fi/bmi_isc_analysis

### Competing Interest Statement

The authors have declared no competing interest.

