## supplemental figure for "Dual states of Bmi1-expressing intestinal stem cells drive epithelial development and tissue regeneration"

**Figure S1. FACS gating scheme for isolating adult or developmental Bmi1-GFP^+^ cell populations**. (A) Adult and (B) E14.5 Bmi1-GFP^+^ epithelial cell populations from murine Bmi1-GFP reporter intestine. Gates from left to right: total cells, singlets, live (propidium iodide negative) CD31/CD45 negative, EpCAM^+^, Bmi1-GFP^+^. Inset: C57Bl/6 (WT) Bmi1-GFP negative control gates.

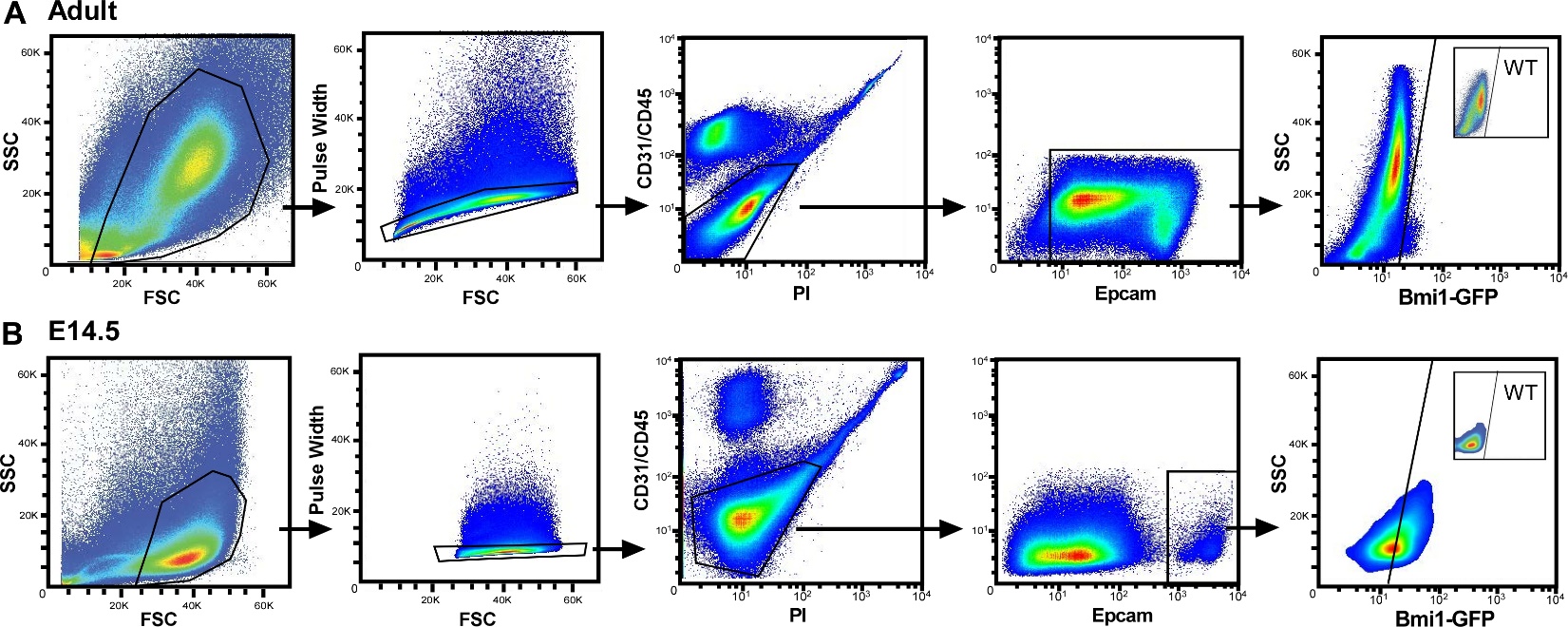

**Figure S2. FACS plots of EpCAM^+^ intestinal cells across developmental time**. C57Bl/6 (WT; top panel), Bmi1-GFP (middle panel) and Lgr5-GFP (bottom panel) at (left to right) E12.5, E14.5, E15.5, E17.5 and Adult.

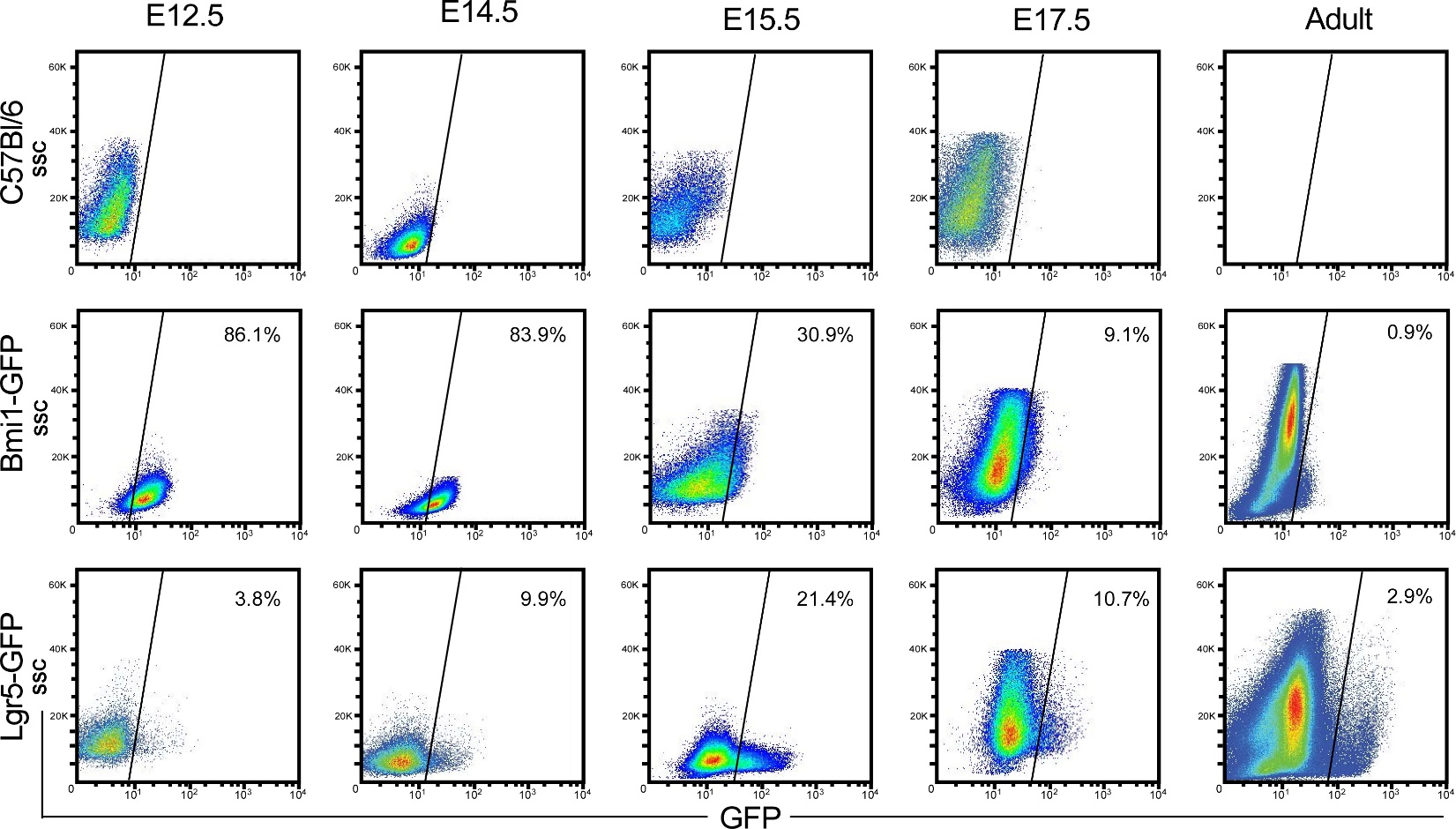

**Figure S3. Lineage tracing from Bmi1- and Lgr5-expressing populations in adult tissues**. (A) Adult intestinal tissue wholemount images lineage traced from Bmi1-TdTomato (TdT, top) or Lgr5-TdT (bottom) at E8.5. Proximal small intestine (PSI), middle small intestine (MSI), distal small intestine (DSI) and Colon. Lineage marked cells express TdTomato (TdT, red). Data are representative of N=11 (Bmi1) and 7 (Lgr5) mice from at least 5, 3 independent experiments. (B) Intestinal tissues from Bmi1 or Lgr5 traced tissues with lineage-derived cells marked by TdT expression (red) and co-stained for EpCAM (white). Top row from Figure 3 with lineage induced at E8.5 and analyzed at E17.5, bottom row with lineage induced at E10.5 and analyzed at E17.5. Arrowheads depict a subset of TdT^+^ progeny. Scale bar = 25 µm. Data are representative of N=3-9 mice for Bmi1 and Lgr5 from at least 2 independent experiments. (C) Induced lineage tracing at E8.5 from Bmi1^+^ cells on an Lgr5-GFP reporter background (Bmi1-Cre^ERT^ x R26R-TdT, Lgr5-GFP), followed by analyses at E17.5. Tissue sections showing Bmi1-derived cell lineage with TdT (red), Lgr5-GFP (green) and stained for EpCAM (white). Arrowheads denote TdT^+^ crypts with Lgr5-GFP^+^ ISCs, scale bar = 25 µm. Data are representative from N=3 mice from 2 independent experiments.

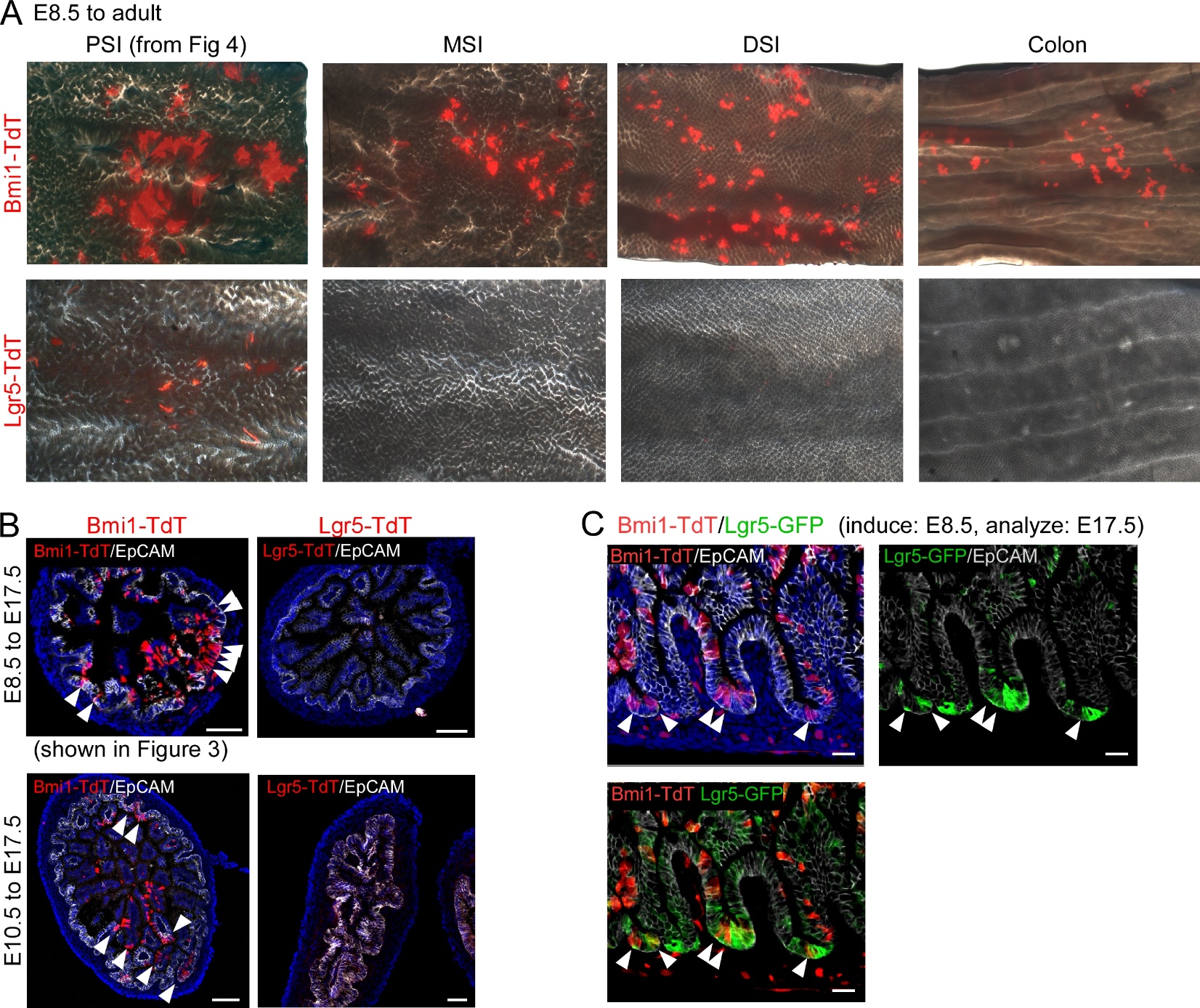

**Figure S4. Clonal analysis of Bmi1 and Lgr5 lineage tracing during development.** Low mag (left) and high mag images of boxed fields (right) of TdTomato-marked clones from Bmi1-cre (top panel) or Lgr5-cre (bottom panel) mice. White arrowheads denote clones, white dashed lines denote the epithelial-mesenchymal boundary Data are representative of N=11, 7 mice from at least 5, 3 independent experiments for Bmi1 and Lgr5 respectively. Related to Figure 3. Scale bar = 25 µm.

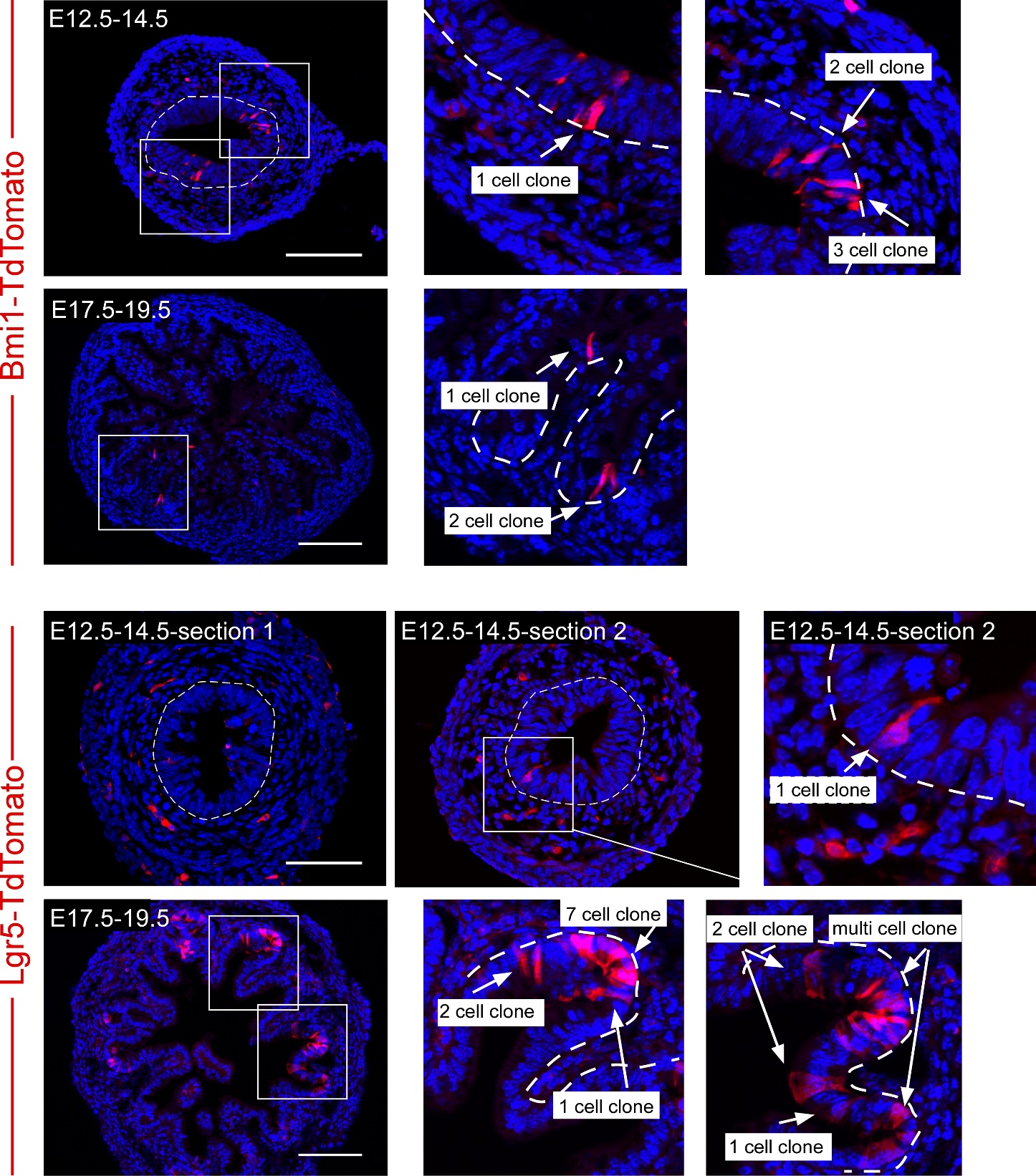

**Sup Fig 5** **scRNA-seq analyses of developmental Bmi1^+^ datasets.** (A) UMAP plot of E12.5 and 17.5 Bmi1^+^ scRNA-seq datasets integrated using the Harmony approach, from Figure 4A, shown here for reference. (B-E) Inference Pseudotime mapped for E12.5 and E17.5 subpopulations and annotated for differentiated lineages. (F) UMAP plots of Bmi1-GFP^+^ cells from E12.5 (top) and E17.5 (bottom) depicting activities of pathways highlighted for Stem/Progenitor, Developmental, and Differentiation. (G) Stem cell populations highlighted in darker color for E12.5 (dark pink) and E17.5 (dark green) on the UMAP from A. (H) Comparison of differentially expressed genes (DEGs) between stem cell clusters from E12.5 and E17.5. DEGs were selected by marker analysis between the identified stem cell populations and other populations in E12.5 or E17.5 independently and then selected using threshold values, logFC > 2 and FDR < 0.01.

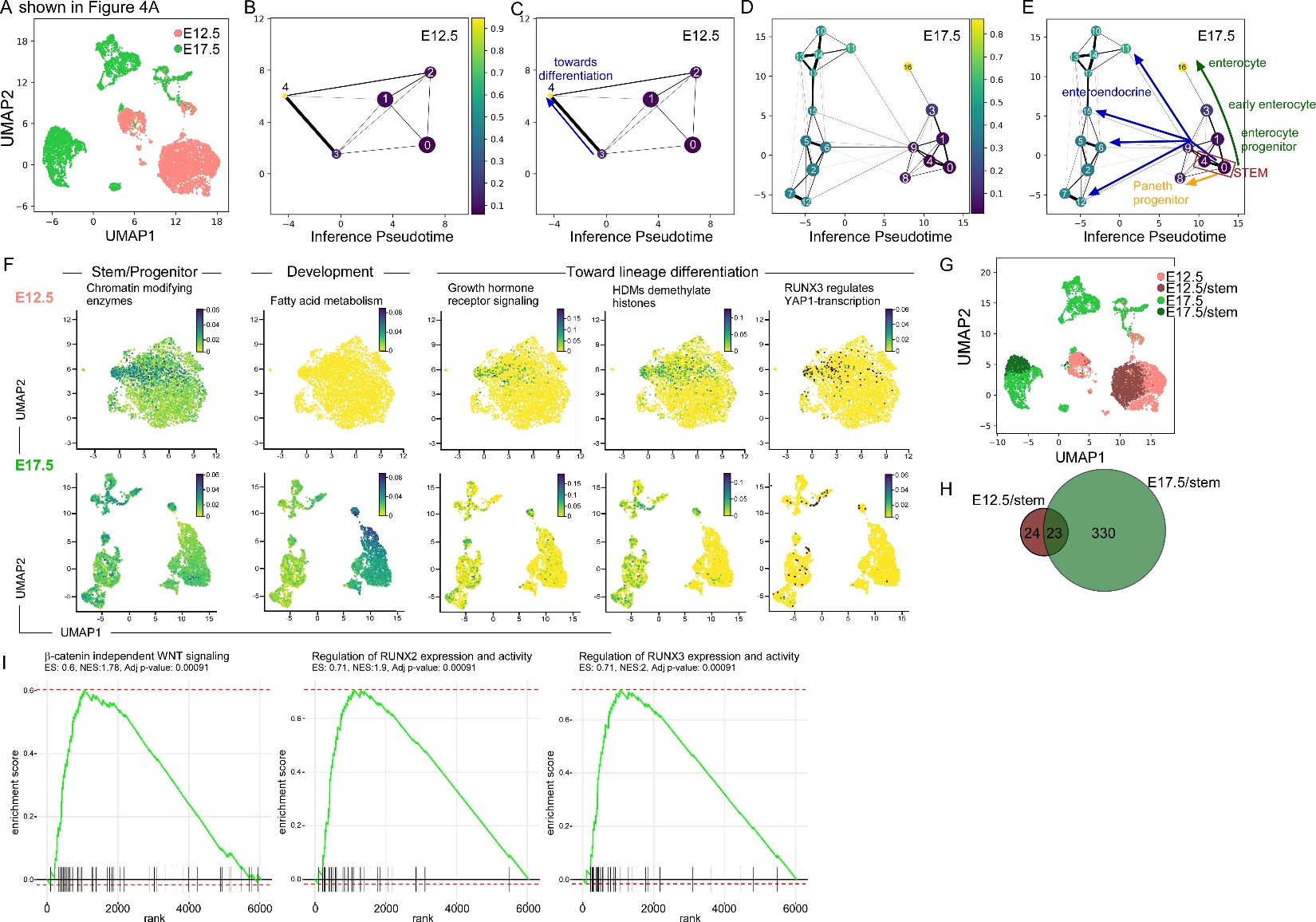

**Figure S6. Bmi1^+^ cells lack enteroendocrine lineage markers in early development.** (A) Intestinal tissue from 48 hr lineage traced progeny from Bmi1 promoter from developmental and adult timepoints, stained with antibodies to EpCAM (white), chromogranin A (Chga, green), and TdTomato traced cells (red). Yellow arrowhead denotes Bmi1-expressing or lineage traced progeny, white arrowhead denotes Bmi1 and Chga double-positive cells, and gray arrowhead denotes cells only expressing Chga. (B) UMAP plot of Bmi1-GFP^+^ scRNA-seq data from Yan et al. (C) Merge of Yan and our Bmi1-GFP^+^ datasets and UMAP visualization depict stem and intestinal subpopulations. (D) Violin plots of stem (*Mki67, Cdk1, Ascl2*) and enteroendocrine cell (EE; *Chga, Chgb*) gene expression and developmental (DEV #) and chromatin regulatory gene (CRG #) scores highlight shared identity of Bmi1-expressing stem population from two different datasets.

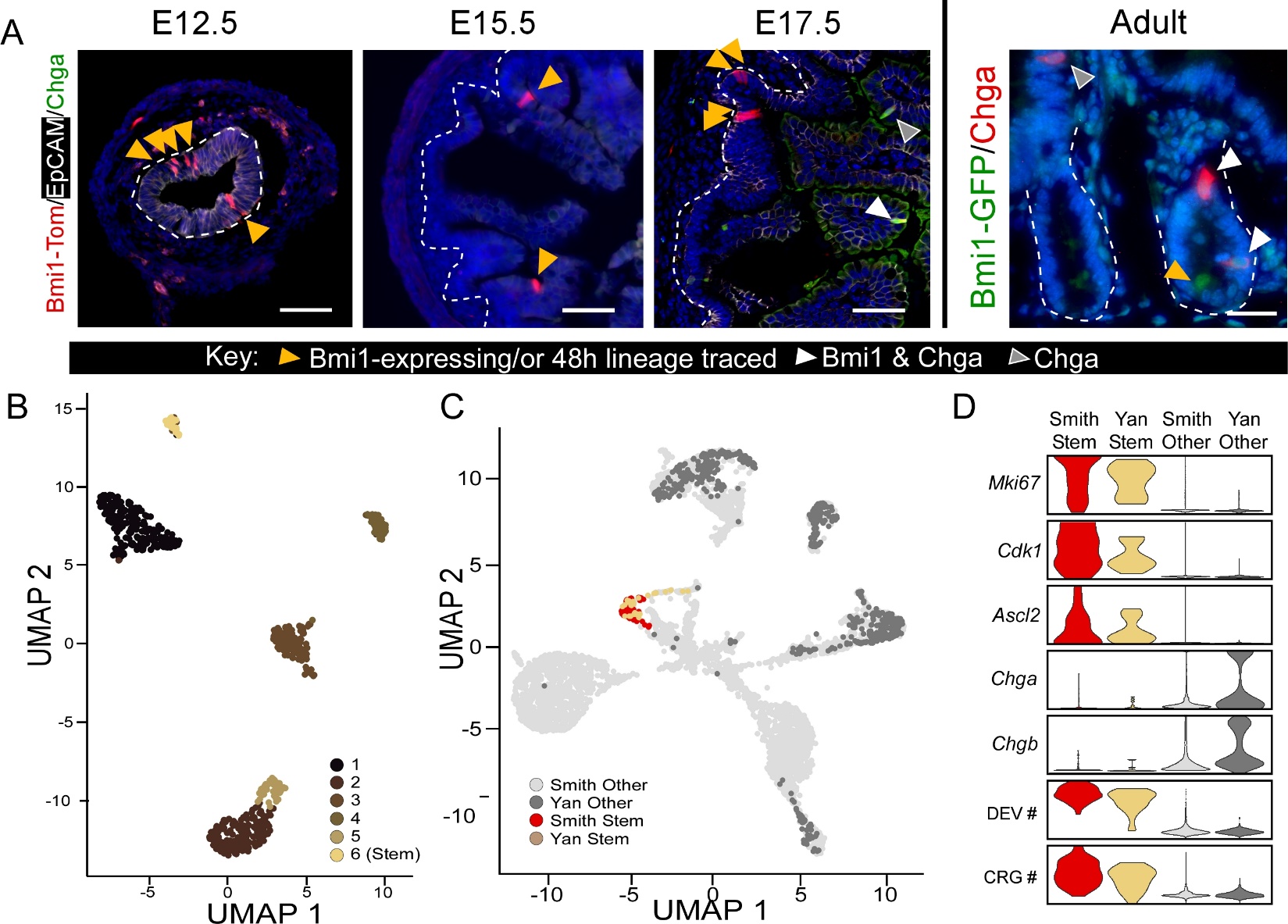

**Figure S7. Characterizing stem and non-stem clusters.** (A) Violin plots of numeral of expressed genes (left) and unique molecular identifiers (right) within each cell, separated by cell type. (B) UMAP plot (left) of adult Bmi1-GFP^+^ cells reveals secretory progenitor (grey) population. We scored this population for genes associated with differentiated enteroendocrine cell (EE) populations (EC, I, D, K, Paneth) and annotated each secretory progenitor cell with their highest scoring cell type (right), confirming this heterogenous progenitor population. (C) Gene expression of magnified UMAP visualization of unique irradiated cluster. The irradiated stem and secretory precursor cell types are differentiated by their expression of ISC genes (*Olfm4, Ascl2, Myc, Cdk4, Cdk6, CD44*). Previous studies have reported post-irradiated emergence of a revival stem cell with unique markers (*Clu, Anxa1, Mif, Npm1, Cxadr*) and a secretory precursor cell (*Chga, Cck, Ghrl, Nts, Neurod1, Serpina1c*) that are recapitulated in our dataset and differentiate this cluster.

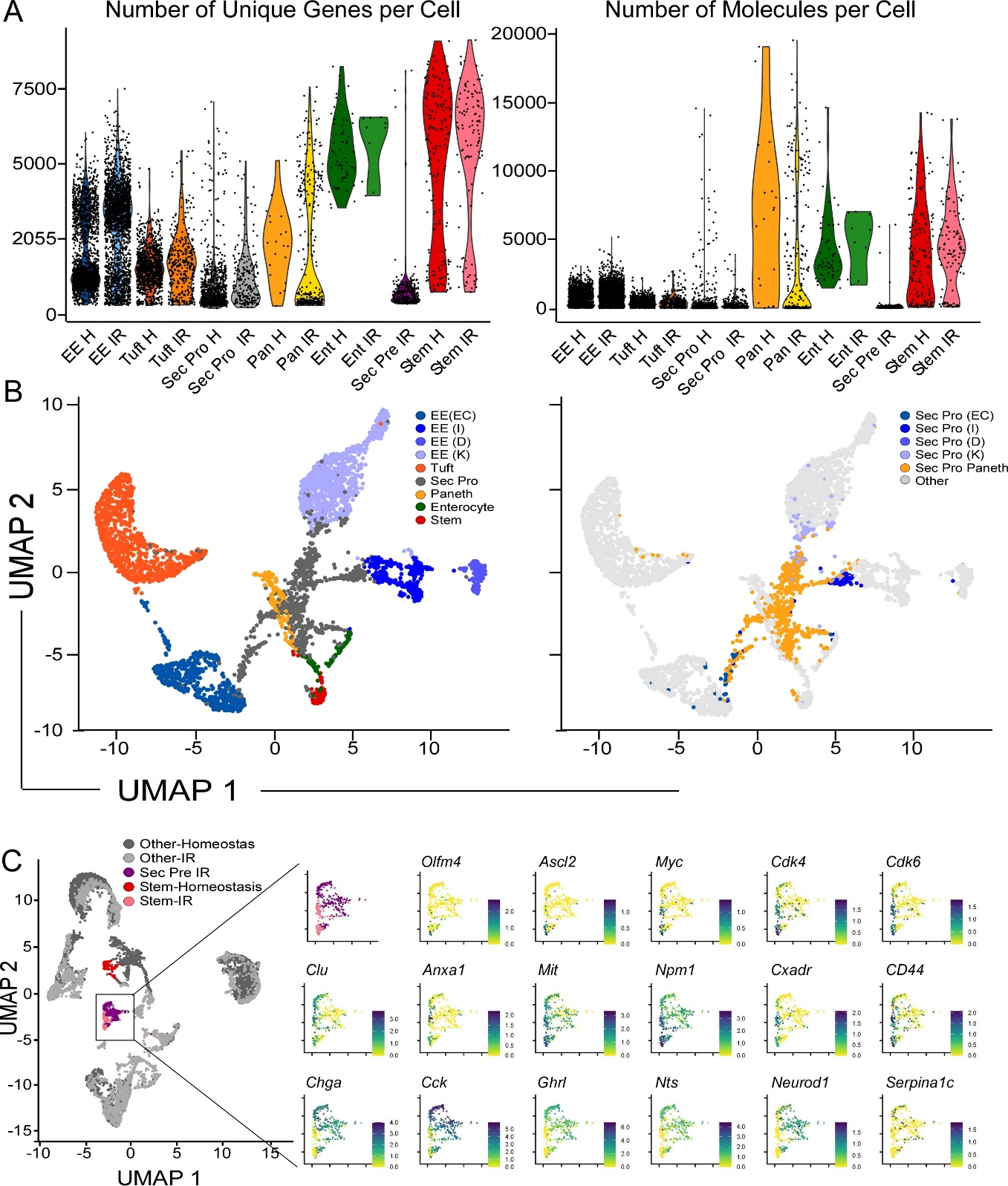
